# Paternal obesity results in placental hypoxia and sex-specific impairments in placental vascularization and offspring metabolic function

**DOI:** 10.1101/2021.03.27.437284

**Authors:** Patrycja A. Jazwiec, Violet S. Patterson, Tatiane A. Ribeiro, Erica Yeo, Katherine M. Kennedy, Paulo C.F. Mathias, Jim J. Petrik, Deborah M. Sloboda

**Author notes:** Authors have equally contributed to the study and are listed in alphabetical order (Patrycja A. Jazwiec, Violet S. Patterson, Tatiane A. Ribeiro). Corresponding Author: D.M. Sloboda McMaster University Department of Biochemistry and Biomedical Sciences 1280 Main St West, Hamilton, Canada.

## Abstract

Paternal obesity predisposes offspring to metabolic dysfunction, but the underlying mechanisms remain unclear. We investigated whether paternal obesity-induced offspring metabolic dysfunction is associated with placental endoplasmic reticulum (ER) stress and impaired vascular development. We determined whether offspring glucose intolerance is fueled by ER stress-mediated changes in fetal hepatic development. Furthermore, we also determined whether paternal obesity may indirectly affect *in utero* development by disrupting maternal metabolic adaptations to pregnancy. Male mice fed a standard chow diet (CON; 17% kcal fat) or high fat diet (PHF; 60% kcal fat) for 8-10 weeks were time-mated with control female mice to generate pregnancies and offspring. Glucose tolerance in pregnant females was evaluated at mid-gestation (embryonic day (E) 14.5) and term gestation (E18.5). At E14.5 and E18.5, fetal liver and placentae were collected, and markers of hypoxia, angiogenesis, endocrine function, and macronutrient transport, and unfolded protein response (UPR) regulators were evaluated to assess ER stress. Young adult offspring glucose tolerance and metabolic parameters were assessed at ∼60 days of age. Paternal obesity did not alter maternal glucose tolerance or placental lactogen in pregnancy but did induce placental hypoxia, ER stress, and altered placental angiogenesis. This effect was most pronounced in placentae associated with female fetuses. Consistent with this, paternal obesity also activated the ATF6 and PERK branches of the UPR in fetal liver and altered hepatic expression of gluconeogenic factors at E18.5. Adult offspring of obese fathers showed glucose intolerance and impaired whole-body energy metabolism, particularly in female offspring. Thus, paternal obesity programs sex-specific adverse placental structural and functional adaptations and alters fetal hepatic development via ER stress-induced pathways. These changes likely underpin metabolic deficits in adult offspring.

**Summary Sentence:** Paternal obesity alters placental vascular structures and is associated with sex-specific compromises in glucose tolerance and metabolism in young offspring

## INTRODUCTION

Obesity prevalence continues to increase globally.^1^ As of 2016, 13% of adults worldwide were classified as obese,^1^ increasing their risk of developing co-morbidities, including coronary heart disease and type 2 diabetes. Obesity risk has traditionally been explained by genetic predisposition and/or lifestyle factors, and the contribution of the early life environment often goes unrecognized.^2^ It is now well established that at least some portion of an individual’s health trajectory is determined early in life, beginning prior to conception (in germ cells) and continuing into the embryonic, fetal, and early postnatal periods.^3^ Indeed, early in development an organism is plastic and responds to environmental signals or factors, changing its developmental trajectory.^3^ One of the most studied early life factors is exposure to maternal obesity. Although the relationship between maternal obesity, offspring obesity, and metabolic dysfunction has been well-established,^4, 5^ whether the paternal lineage shares a similar contribution, has been largely overlooked and understudied.

Some, albeit few, clinical data exist demonstrating a link between paternal diet and/or obesity, and offspring obesity risk and metabolic dysfunction.^6^ In experimental models, paternal diet-induced obesity results in impaired offspring glucose regulation^7, 8^ and altered mitochondrial function,^9^ but which specific metabolic organs/pathways are impacted and when, remains unclear. Like maternal obesity, paternal obesity impacts offspring in a sex-specific manner, as females appear more vulnerable to paternal-lineage induced early life adversity. In mice, female embryos derived from obese males not only have delayed development, but far fewer of them reach the blastocyst stage.^10^ Most offspring impairments have been attributed to the damaging effects of paternal obesity on sperm^11, 12, 13^ and on sperm DNA methylation,^14^ chromatin structure, and non-coding RNAs.^8, 15^ But despite the fact that the sperm epigenome also regulates growth and development of the placenta,^16^ few studies have investigated the impact of paternal obesity on placental development,^17, 18, 19^ and none have investigated whether these placental changes alter maternal metabolic adaptation to pregnancy. As a result, whether paternal obesity channels indirect effects on the developing fetus/offspring through either maternal metabolic impairments, or through the placenta, is virtually unknown.

The placenta is a highly vascularized interface between the developing organism and its mother, providing a continuous supply of oxygen and nutrients to sustain fetal growth and development.^20^ Although we know that maternal obesity impairs placental vascularization,^4, 21^ induces hypoxia and oxidative stress,^4, 22^ endoplasmic reticulum (ER) stress,^23^ and alters fetal metabolic development,^24^ little to no data exist regarding the impacts of paternal obesity on the placenta, or on maternal or fetal metabolic development. Here, we investigated the impacts of paternal obesity on maternal glucose metabolism, placental vascular development, and placental ER stress and activation of the unfolded protein response (UPR). We show that despite having no effect on maternal glucose metabolism, paternal obesity induced placental hypoxia, altered placental angiogenesis, and altered placental insulin signaling without impacting macronutrient transport. We also show that at least two arms of the UPR were activated (PERK and IREα) in placentae of fetuses sired by obese males. These changes were more prevalent in placentae associated with female fetuses. Since placental function largely determines fetal development, we also determined whether paternal obesity-induced impairments in the placenta were associated with UPR activation and ER stress in a major fetal metabolic organ, the liver. We found that the UPR was activated in the liver of fetuses of obese fathers and was associated with increased expression of gluconeogenic transcriptional activators. Young adult offspring (particularly females) born to obese fathers were glucose intolerant. We speculate that these early (fetal) changes in metabolic control could partly account for our observed changes in offspring whole body energy metabolism. Our data suggest that although both sexes have postnatal impaired metabolic activity, females are particularly vulnerable to paternal-lineage obesity, and that this may be due in part, to impairments in placental oxygenation.

## METHODS

### Animal Model

#### Diet-Induced Obesity in Male Mice

All animal experiments were approved by the Animal Research Ethics Board at McMaster University (Animal Utilization Protocol 16-09-35) and were in accordance with the Canadian Council on Animal Care guidelines. We used a model of high fat (HF) diet-induced obesity in C57BL/6J male mice (Supplemental Fig. S1). Six-week-old male mice were randomized to either a: 1) *Control (CON) group*: fed a standard chow diet (n = 49; 17% kcal fat, 54% kcal carbohydrates, 29% kcal protein, 3 kcal/g; Harlan 8640 Teklad 22/5 Rodent Diet) or a 2) *Paternal High Fat (PHF) group*: fed a HF diet (n = 61, 20% kcal protein, 20% kcal carbohydrates, 60% kcal fat, 5.21 kcal/g, Research Diets Inc. D12492) *ad libitum* with free access to water for 8-10 weeks. All male mice were housed in the same room maintained at 25°C and a 12-hour (hr) light, 12-hr dark cycle. Food intake and body mass were measured weekly. Body adiposity was assessed in CON and PHF male mice using a body composition analyzer (Bruker Minispec LF90-II) prior to diet randomization (baseline or 0 weeks), and 5 and 7 weeks after consuming their respective diets. To assess glucose tolerance, a standard intraperitoneal (*i.p.*) glucose tolerance test (GTT) was performed on 12hr fasted male mice at baseline and after 5 weeks of diet consumption. Briefly, CON and PHF male mice were injected with glucose (G5767; Sigma-Aldrich; 2 g/kg, *i.p.*). Blood glucose was repeatedly measured through tail vein sampling using Accu-Check Aviva blood glucometer (Roche Diagnostics) prior to glucose injection (0) and at 20, 30, 40, 60, 90, and 120 minutes (mins) after glucose injection.^25^

#### Timed Mating and Pregnancy

After 8-10 weeks of CON or PHF diet, male mice were timed-mated with virgin C57BL/6J female mice to generate pregnancies. One or two female mice were housed in the same cage overnight with a male mouse. Mating was confirmed by the presence of a copulation plug the following morning and designated at embryonic day (E) 0.5. Pregnant female mice were housed individually and fed a standard chow diet (17% kcal fat, 54% kcal carbohydrates, 29% kcal protein, 3 kcal/g; Harlan 8640 Teklad 22/5 Rodent Diet) *ad libitum* and provided free access to water throughout gestation. Maternal food intake and body mass was measured throughout gestation. Pregnant mice either underwent glucose tolerance testing (see below) or were sacrificed at mid-gestation (E14.5; n = 10) and term gestation (E18.5; n = 10-13) by cervical dislocation. Placentae and fetal liver samples were collected from one randomly selected male and one female fetus in each litter at E14.5 and E18.5 (see below). Fetal tails were collected to sex fetoplacental units. Briefly, DNA was extracted from fetal tail, *Sry* was PCR-amplified using 5’ TTGTCTAGAGAGCATGGAGGGCCATGTC 3’ and 5’CCACTCCTCTGTGACACTTTAGCCCTCCG 3’ primers.

For offspring studies, a separate cohort of pregnant female mice was allowed to deliver and at birth litters were standardized six pups (preferentially to three male pups and three female pups) to ensure standardized nutrition during lactation. Offspring food intake and body mass were measured weekly. Male and female CON and PHF offspring were subjected to metabolic assessments (see below) as young adults at postnatal day 53 (P53) and P60, after which they were sacrificed by cervical dislocation at P67.

#### Maternal Glucose Metabolism Assessments

To test whether paternal obesity impacted maternal metabolic adaptation to pregnancy, pregnant females were subjected to a GTT. At mid-gestation (E14.5), dams mated with CON and PHF males (n = 10) were fasted for 6hrs and subjected to a standard intraperitoneal *(i.p.)* GTT with glucose (G5767; Sigma-Aldrich; 1.5g/kg, *i.p.*). In another cohort at term gestation (E18.5), dams mated with CON and PHF males (n = 10-13) were fasted for 6hrs and subjected to a standard oral GTT. Due to the difficulty of successfully completing an *i.p.* injection with a full gravid uterus, pregnant dams were gavaged with glucose (G5767; Sigma-Aldrich; 2g/kg). At E14.5 and E18.5, maternal blood glucose was repeatedly measured through tail vein sampling using an Accu-Check Aviva blood glucometer (Roche Diagnostics) prior to glucose administration (0) and 15, 20, 30, 40, 60, 90, and 120mins after glucose administration.

#### Offspring Metabolic Assessments

Metabolic profiling was performed on male and female CON (n = 14/sex) and PHF (n = 8/sex) offspring using Comprehensive Lab Animal Monitoring System (CLAMS; Columbus Instruments) at postnatal day (P) 53.^26^ Briefly, offspring were placed into individual CLAMS cages at 15:00hrs and acclimatized for 24hrs prior to acquiring measurements. Measurements included: food consumption, water consumption, total horizontal motor activity, heat production, oxygen consumption (VO_2_), carbon dioxide production (VCO_2_), respiratory exchange ratio (RER), carbohydrate oxidation (carbOx), and lipid oxidation (lipOx). Measurements were acquired every 20mins for 48hrs.

At P60 a standard *i.p.* GTT was performed on fasted male and female CON (n = 6/sex) and PHF (n = 7-8/sex) offspring. Offspring were injected with glucose (G5767; Sigma-Aldrich; 2g/kg, *i.p.*) and blood glucose was repeatedly measured through tail vein sampling using an Accu-Check Aviva blood glucometer (Roche Diagnostics) prior to glucose injection (0) and 20, 30, 40, 60, 90, and 120mins after glucose injection.

#### Insulin ELISAs

All serum insulin concentrations were quantified using a commercially available insulin immunoassay kit (32380; Toronto Bioscience) as per manufacturer’s instructions using Synergy H4 Hybrid microplate reader (BioTek Instruments). Insulin concentrations were interpolated from a standard curve determined by curve of 4-parameter as per manufacturer’s instructions. Homeostatic model assessment of insulin resistance (HOMA-IR) was calculated by multiplying blood glucose concentrations (mmol/L) by serum insulin concentrations (mU/mL), and dividing this value by 22.5.^27^

### Molecular Analyses

#### RNA Extraction

Placentae and livers from one male and one female fetus were collected from each pregnancy, cut in half, snap frozen in liquid nitrogen, and stored at −80°C until RNA extraction. Placental and hepatic tissues were homogenized in 900μl of TRIzol reagent (15596018; Invitrogen) using glass homogenizing beads (10064583; ACROS Organics) and a homogenizer (MP116004500; MP Biomedicals). Homogenates were centrifuged at 12,000*g* for 10mins at 4°C. Supernatant was removed, added to 300μl of chloroform (BP1145-1; FisherBioReagents), and thoroughly mixed. Samples were incubated at room temperature (RT) for 3mins, and centrifuged 12,000*g* for 10mins at 4°C. The aqueous layer was removed, mixed with 500μl of isopropanol (BP26181; FisherBioReagents), incubated at RT for 20mins, and then centrifuged at 12,000*g* for 10mins at 4°C. Supernatant was removed, the remaining RNA pellets were washed twice in 75% ethanol (EtOH), and then reconstituted in 20μl of ultrapure water (UP-H_2_O). RNA was quantified using a NanoDrop 2000 spectrophotometer (ThermoScientific) with the NanoDrop 2000/2000c software (ThermoScientific). RNA quality was determined using ratio of absorbance at 260nm:280nm (A_260_/A_280_) and 260nm:230nm (A_260_/A_230_). RNA samples with an A_260_/A_280_ and A_260_/A_230_ ratio of >2.0 and >1.5, respectively, were used to generate complementary (cDNA). RNA samples were stored at −80°C until required for cDNA synthesis.

#### Complementary DNA Synthesis

Two micrograms of placental and hepatic RNA were used for first strand cDNA (cDNA) synthesis using SuperScript IV VILO Master Mix with ezDNase enzyme (11766050; Invitrogen) as per the manufacturer’s instructions. Complementary DNA samples were diluted to 1:100 in UP-H_2_O and stored at −80°C until required for quantitative PCR (qPCR) assays.

#### Quantitative PCR Assays

Quantitative PCR assays were performed as before.^4^ Primer sets (Supplemental Table S1) of candidate genes were designed using Primer Basic Local Alignment Search Tool (Primer-BLAST) software available at the National Center for Biotechnology Information (https://www.ncbi.nlm.nih.gov/tools/primer-blast/, NCBI). Primer conditions were adjusted to the following cycling conditions: PCR Product Size: min: 50bp, max: 150bp; Primer Melting Temperatures (T_m_): min: 58°C, opt: 60°C, max: 62°C; Max T_m_ Difference: 2°C. Primer pairs were designed to span exon-exon junctions. Primers were manufactured by Invitrogen (Invitrogen Life Technologies).

The LightCycler 480 SYBR Green I Master and the LightCycler 480 system (Roche Diagnostics) were used for quantifying transcript levels. Each reaction consisted of 2.5μL of cDNA (1:100 dilution), 1μL of 5μM forward primer and 1μL of 5μM reverse primer for the gene of interest, 0.5μL of UP-H2O, and 5μL of Lightcycler 480 SYBR Green I Master (04887352001; Roche Diagnostics). The cycling conditions were: enzyme activation at 95°C for 5mins, amplification of the gene product through 60 successive cycles of 95°C for 10sec, 60°C for 10sec, and 72°C for 10sec, and melting curve beginning at 65°C and ending at 95°C. Each qPCR assay for each gene of interest contained a standard curve (10-fold serial dilution of pooled cDNA generated by pooling equal volumes of cDNA from each sample), cDNA of samples, and a non-template control (UP-H_2_O). Transcript levels were quantified in triplicate for each standard and sample. Gene expression data were normalized to the geometric mean of at least two housekeeper genes (Supplemental Table S2).

### Placental Histology

#### Quantification of Placental Trophoblast Giant Cells

To determine whether paternal obesity altered placental endocrine-mediated maternal adaptations to pregnancy, a subset of placental samples dissected at E14.5 were processed for the quantification of placental giant cells as a marker of placental lactogen production. These cells are responsible for the secretion of placental lactogen,^28^ which mediates maternal glucose metabolism. Placental samples were immersion-fixed in modified Davidson’s Fixative for 8hrs at RT prior to being processed and paraffin-embedded. Placentae were serially sectioned at 8μm. Tissue sections were deparaffinized (Histoclear, Electron Microscopy Sciences) and rehydrated in a graded EtOH series. Three replicate placental sections 80μm apart were stained for polysaccharides (including glycogen) using Periodic Acid Schiff (PAS, 395B-1KT, Sigma-Aldrich) (n = 7-11/group/sex). Parietal trophoblast giant cells were counted using Nikon NIS Imaging Analysis Software (v.5.20.02).

#### Placental Immunostaining

To quantify angiogenic markers and hypoxia, a subset of E14.5 and E18.5 placentae were immersion-fixed in modified Davidson’s Fixative for 8hrs at RT or in 4% paraformaldehyde (50980486; FisherScientific) for 24hrs at RT, respectively. Fixed tissues were paraffin-embedded and serially sectioned at 4μm. Three replicate sections from each placental sample were immunolabelled for: hypoxia marker carbonic anhydrase [CA IX]), pro-angiogenic factors (vascular endothelial growth factor A [VEGF-A], vascular endothelial growth factor receptor 2 [VEGFR-2]), endothelial cell marker (cluster of differentiation 31 [CD31]), and pericyte marker (α-smooth muscle actin [α-SMA])(Supplemental Table S3). Endogenous peroxidases were inhibited with 30%v/v hydrogen peroxide (H_2_O_2_; diluted in TBS; H324-500; Fisher Scientific). Antigen retrieval was performed with 10mM sodium citrate buffer with Tween at 95°C for 12mins (pH 6.0, washed 2×5mins in TBS). Non-specific binding was blocked with 5% bovine serum albumin (BSA; in TBST pH 7.4, A2153; Sigma-Aldrich) for 10mins at RT Sections were then incubated with primary antibody (diluted in 1% BSA/TBST) in a humidified chamber overnight at 4°C. Sections were then incubated with a biotinylated secondary antibody (1:100 dilution in TBST, PK6101; Vector Laboratories) or fluorescently-conjugated secondary antibodies (Sigma-Aldrich, Canada and Vector Laboratories) for 1hr. For chromogenic immunohistochemistry, sections were incubated with an avidin-biotin peroxidase complex following VectaStain ABC kit protocols (PK-4000; Vector Laboratories) for 1hr. Biotinylated antibody was visualized with chromagen development using 3,3’ diaminobenzidine (DAB) peroxidase (SK-4100; Vector Laboratories), then sections were counterstained with Mayer’s Hematoxylin (MHS32; Sigma-Aldrich), and mounted with coverslips using Permount Mounting Media (SP15500; FisherScientific).

The proportion of positive immunostaining relative to placental area was determined using a threshold for DAB-positive staining or the proportion of fluorescent-positive cells using NIS Elements Software (Nikon Instruments Inc). Image analysis was performed using an Olympus BX-61 microscope and integrated morphometry software (MetaMorph) (CA IX, VEGF-A, α SMA) or on a Nikon Eclipse NI microscope and Nikon NIS Elements Imaging Software (v.4.30.02) (CD31). The ratio of CD31 to α-SMA immunopositive area was used as a marker of vessel maturity. All image analyses were performed by an investigator blinded to the study groups.

### Immunoblotting

#### Total Protein Extraction

Placental and fetal hepatic tissues were homogenized in 300μl and 200μl of extraction buffer (50mM HEPES, 150mM NaCl, 100mM NaF, 10mM sodium pyrophosphate, 5mM EDTA, 250mM sucrose, 1% Triton-X, 1mM sodium orthovanadate, 1% protease inhibitor tablet) using ceramic homogenizing beads (19-646-3; Omni International) to extract total protein. Samples were centrifuged at 14,000*g* for 15mins at 4°C. Total protein concentration was quantified using the Pierce BCA Protein Assay Kit (23225; ThermoFisher Scientific) according to the manufacturer’s protocol.

#### Nuclear Protein Extraction

Nuclear protein was extracted from E14.5 CON and PHF placentae. Briefly, tissue was homogenized in 500μl of hypotonic lysis buffer (10mM HEPES, pH 7.9, 1.5mM MgCl_2_•6 H_2_O, 10mM KCl, 1% protease inhibitor tablet) using ceramic homogenizing beads (19-646-3, Omni International), centrifuged at 11,000*g* for 20mins, and the supernatant was discarded (cytoplasmic fraction). The pellet (nuclei) was resuspended in 70μl of extraction buffer (20mM HEPES, pH 7.9, 1.5mM MgCl_2_•6 H_2_O, 420mM NaCl, S671-3, 0.2mM EDTA, 25%v/v glycerol, 1% protease inhibitor tablet) and homogenized again as described above. Homogenates were incubated at RT for 30mins, centrifuged at 21,000*g* for 5mins, and nuclear fractions were quantified using the Pierce BCA Protein Assay Kit (23225, ThermoFisher Scientific) as per manufacturer’s protocol.

#### Protein Detection and Quantification

Total and nuclear protein was separated using gel electrophoresis (7.5%-15% separating gel). Proteins were transferred to membranes (1620177, BioRad) using a TransBlot^®^ Turbo^TM^ Transfer System (1704150, BioRad). Blots were then blocked in 5% BSA (in TBST, A2153-1KG, Sigma-Aldrich) for 1hr. Blots were incubated overnight in rabbit anti-mouse primary antibody for proteins of interest (Supplemental Table S4) and with horseradish peroxidase-conjugated goat anti-rabbit IgG secondary antibodies (ab6721, Abcam). Proteins of interest were detected by incubating blots in Clarity^TM^ Western ECL Blotting Substrate (1705061; BioRad) or Clarity Max^TM^ Western ECL Blotting Substrate (1705062; BioRad) and images were captured using a ChemiDoc MP Imaging System (1708280; BioRad). Densitometric quantification was completed using ImageLab software (ImageLab Software for PC Version 6.0.1 SOFT-LIT-170-9690-ILSPC; Bio-Rad). All protein data is expressed as a relative to concentration of an internal control as fold change.

#### Phosphorylated Protein Detection and Quantification

Phosphorylated proteins were separated using SDS-PAGE (7.5%-15% separating gel). Proteins were transferred to membranes (1620177, BioRad) using a TransBlot^®^ Turbo^TM^ Transfer System (1704150, BioRad). Blots were then blocked in 5% BSA (in TBST, A2153-1KG, Sigma-Aldrich) for 1hr. Blots were incubated overnight in rabbit anti-mouse primary antibody for phosphorylated protein of interest (Supplemental Table S4), then incubated in a horseradish peroxidase-conjugated goat anti-rabbit IgG secondary antibody (ab6721, Abcam). Phosphorylated proteins of interest were detected by incubating blots in Clarity Western ECL Blotting Substrate (1705061; BioRad). Images were captured using a ChemiDoc MP Imaging System (1708280; BioRad). Blots were incubated in stripping buffer (21059, ThermoFisher Scientific) for 5mins, washed twice in TBST for 15mins, then blocked in 5% BSA (in TBST pH 7.4, A2153; Sigma-Aldrich) for 1hr. Blots were probed with the rabbit anti-mouse primary antibody binding to the total protein (phosphorylated and unphosphorylated; Supplemental Table S4) protein of interest. Total protein was detected by incubating blots in Clarity Western ECL Blotting Substrate (1705061; BioRad). Images were captured using a ChemiDoc MP Imaging System (1708280; BioRad).

### Statistical Analyses

For all pregnancy and offspring analyses, the sample size represents the number of litters analyzed, where a representative male and female were analyzed from a pregnancy (or litter) that was sired by either a CON or PHF male mouse. Therefore, each placental and hepatic sample and each offspring measure was sired by an individual male and represents a replicate of one. All data (except for metabolic activity in offspring assessed using CLAMS) were analyzed either by a Student’s t test, repeated measures two-way ANOVA or mixed-effects model with paternal diet and time as factors, or two-way ANOVA with paternal diet and fetoplacental sex as factors. Bonferroni’s post-hoc was used for multiple comparisons where appropriate. Data that were not normally distributed or with unequal variance were analyzed using a Mann-Whitney U test or Student’s t test with Welch’s correction, respectively. All data are presented as mean ± standard error of the mean (SEM) unless otherwise indicated. In all cases, significance (*) was set at *P* < 0.05. Data were analyzed using GraphPad Prism (GraphPad Prism 6.01 for Windows and GraphPad Prism 8.4.2 for MacOS, GraphPad Software, La Jolla California USA, www.graphpad.com).

Offspring metabolic activity (CLAMS) data were analyzed using linear mixed-effects model (LMM) using the lme4 package (R package, RRID:SCR_015654) with the main effects and multiple comparisons determined by Satterthwaite’s method of approximation in lmerTest (R package, RRID:SCR_015656). Response variable data were separated into light (7:00 to 19:00) and dark (19:00 to 7:00), and average values were taken for heat production, VO_2,_ VCO_2_, RER, carbOx, and lipOx. Sums were taken for food consumption, water consumption and activity. Linear mixed-effects model was performed with time period, mouse sex, and paternal diet as fixed variables, and mouse ID and litter as nested random effects.

## RESULTS

### Male mice fed a HF diet are obese, glucose intolerant, and insulin resistant

At baseline prior to diet feeding, fasting blood glucose and glucose tolerance were similar between sires randomized to standard chow (CON) or HF diet (Supplemental Fig. S2A-C). Consistent with previous work,^25^ male mice consuming a HF diet became obese, characterized by increased body mass and body adiposity compared to CON male mice (Supplemental Fig. S2D,E). After five weeks of consuming a HF diet, PHF mice were hyperglycemic, glucose intolerant, hyperinsulinemic, and insulin resistant (Supplemental Fig. S2F-J). Therefore, PHF males were metabolically compromised prior to mating.

### Paternal obesity alters placental angiogenesis and results in placental hypoxia

Angiogenesis is critical for the development of a placental vascular network to support function; processes that are regulated by hypoxia.^29^ We quantified the proportion of positive immunostaining of hypoxia marker CA IX in placentae derived from CON and PHF males. We observed that CA IX immunostaining was increased in PHF placentae compared to CON placentae (P_Diet_ = 0.0003) in male (P = 0.0174; Fig. 1A) and female (P = 0.0077; Fig. 1A) placentae at E14.5, and this difference persisted to E18.5 (P_Diet_ < 0.0001, P_Sex_ < 0.0001) in PHF male (P < 0.0001; Fig. 1B) and female (P < 0.0001; Fig. 1B) placentae compared to CON. Consistent with this, we found increased protein levels of HIF-1α in PHF placentae at E14.5 (CON male; 1.0328 ± 0.091, CON female; 1.320 ± 0.321, PHF male; 3.548 ± 1.293, PHF female; 2.566 ± 0.718; P_Diet_ = 0.0207)). HIF-1α was also increased at E18.5, but only in female placentae (CON male; 0.932 ± 0.179, CON female; 1.172 ± 0.138, PHF male; 0.750 ± 0.192, PHF female; 1.309 ± 0.122; P_Sex_ = 0.0208).

**Figure 1.**
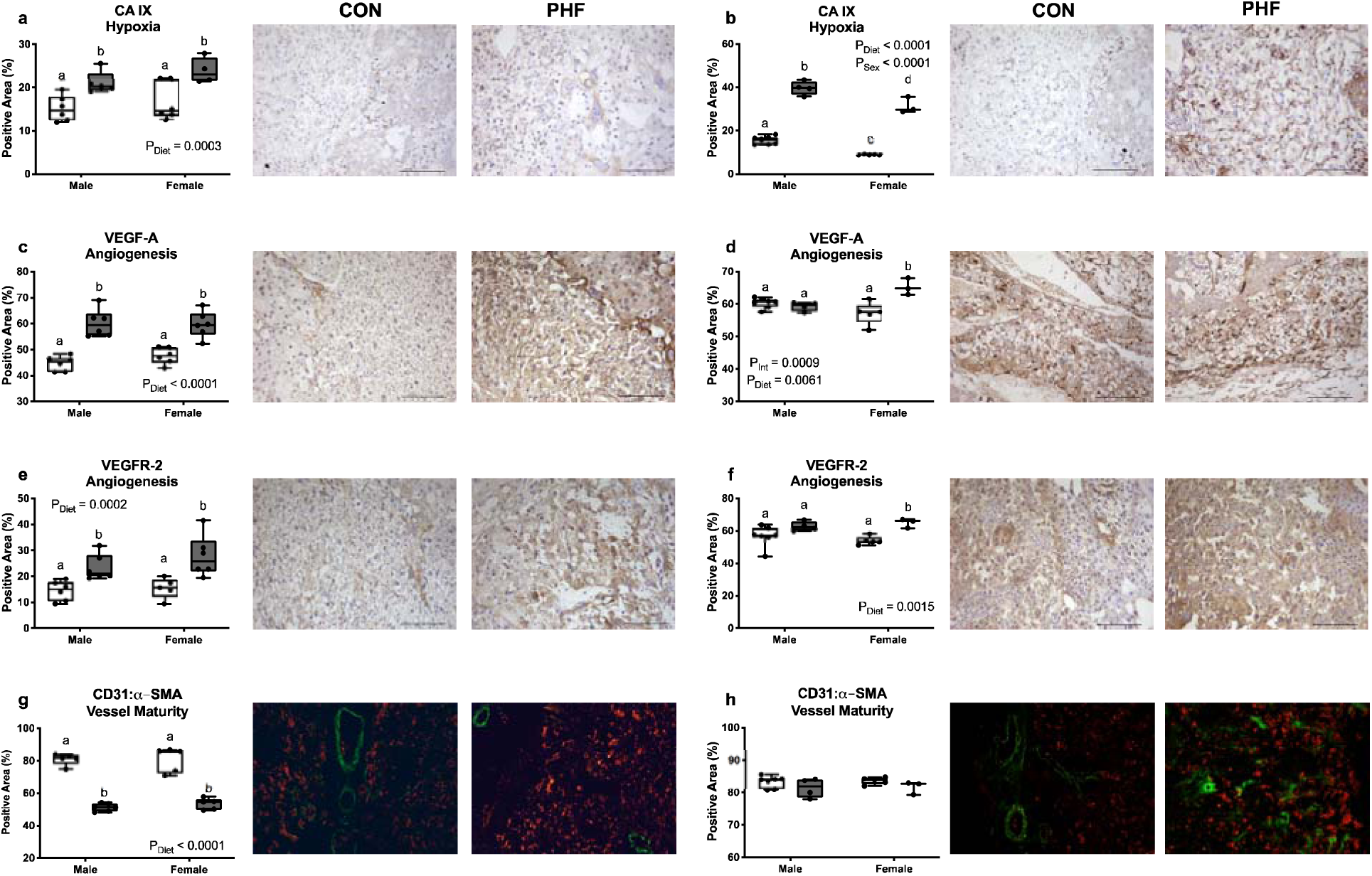
Paternal obesity results in hypoxic placentae with increased number of immature blood vessels. Male and female CON and PHF placentae were immunostained for carbonic anhydrase (CA IX, hypoxia marker), vascular endothelial growth factor A (VEGF-A, proangiogenic factor), vascular endothelial growth factor receptor 2 (VEGFR-2, proangiogenic factor), cluster of differentiation 31 (CD31, endothelial cell marker), and α-smooth muscle actin (α-SMA, pericyte marker). Box plots represent proportions (%) of immunopositive staining and representative images are from female placental samples. Male placentae showed similar staining patterns. (**a**) Percent immunopositive area of CA IX staining in E14.5 and (**b**) E18.5 placentae, (**c**) VEGF-A staining in E14.5 and (**d**) E18.5 placentae, (**e**) VEGFR-2 staining in E14.5 and (**f**) E18.5 placentae, and (**g**) CD31:α-SMA in E14.5 and (**h**) E18.5 placentae with representative images (Scale Bars = 100μm). CON; n = 5-7, PHF; n = 3-6. Data are presented as box and whisker plots; min to max with a line representing the median. Two-way ANOVA with main effects of paternal diet and placental sex as factors using Bonferroni’s post-hoc for multiple comparison. Box plots with different letters indicate significance *P* < 0.05. CON = Control (open box plots); PHF = Paternal high fat diet-induced obesity (grey box plots).

Since we found that PHF mid-gestation and term gestation placentae appeared hypoxic, we next determined whether paternal obesity impacted placental angiogenesis. We measured the proportion of positive immunostaining of pro-angiogenic factor VEGF-A and its receptor VEGFR-2, endothelial cell marker CD31, and pericyte marker α-SMA. We used CD31:α-SMA immunostaining ratio as a marker of vessel maturity. The proportion of positive VEGF-A placental immunostaining was increased in male (P_Diet_ < 0.0001, P < 0.0001) and female PHF placentae (P = 0.0002; Fig. 1C), although at E18.5 this difference persisted only in PHF female placentae (P_Int_ = 0.0009, P_Diet_ = 0.0061,; P = 0.0014; Fig. 1D). These patterns of change were consistent with an increased proportion of VEGFR-2 immunostaining in PHF male (P_Diet_ = 0.0002, P = 0.0209; Fig. 1E) and female placentae (P = 0.0029; Fig. 1E) at E14.5, and in female (but not male) PHF placentae at E18.5 (P_Diet_ = 0.0015, P = 0.0096; Fig. 1F). At E14.5, the ratio of CD31:α-SMA immunostaining was lower (P_Diet_ < 0.0001) in both male and female PHF placentae (P < 0.0001; Fig. 1G) compared to CON, but at E18.5 was similar between groups (P_Diet_ = 0.0575; Fig. 1H). Collectively, these data indicate that placentae derived from PHF sires have increased angiogenesis but produce immature blood vessels.

Decreased vessel maturity is supported by our observations that paternal obesity reduced the expression levels of transcription factors that mediate blood vessel development. Heme oxygenase (HMOX1) facilitates blood vessel development through regulating matrix metalloproteinases MMP14 and MMP2.^30^ We found that paternal obesity reduced transcript levels of *Hmox1* in both male and female placentae at E14.5 (CON male; 0.807 ± 0.058, CON female; 0.892 ± 0.099, PHF male; 0.644 ± 0.025, PHF female; 0.708 ± 0.069; P_Diet_ = 0.0203), but not at E18.5 (CON male; 1.660 ± 0.328, CON female; 1.629 ± 0.225, PHF male; 1.415 ± 0.255, PHF female; 1.407 ± 0.295). *Mmp14* transcript levels were decreased in PHF female placentae at E14.5 (CON male; 0.917 ± 0.055, CON female; 0.852 ± 0.028, PHF male; 0.758 ± 0.075, PHF female; 0.612 ± 0.045; P_Diet_ = 0.0014, P = 0.0152), but these effects did not persist to E18.5 (CON male; 0.743 ± 0.137, CON female; 0.817 ± 0.061, PHF male; 0.893 ± 0.131, PHF female; 0.947 ± 0.164). *Mmp2* transcript levels were unchanged at E14.5 (CON male; 0.901 ± 0.110, CON female; 0.860 ± 0.091, PHF male; 0.604 ± 0.060, PHF female; 0.806 ± 0.080; P_Diet_ = 0.0580) and at E18.5 (CON male; 0.753 ± 0.132, CON female; 0.748 ± 0.072, PHF male; 0.936 ± 0.150, PHF female; 0.949 ± 0.219). Therefore, it appears that paternal obesity may interfere with placental development by altering factors associated with blood vessel development and cellular matrix integrity.

### Paternal obesity activates PERK and IRE1**α** branches of the UPR in mid-gestation and term gestation placentae

Placental hypoxia and angiogenic processes have been associated with ER stress,^31, 32^ so we therefore set out to determine whether paternal obesity-induced disruptions in placental angiogenesis were associated with ER stress. We quantified protein levels of the ER stress chaperone GRP78, as well as regulators and effectors of three branches of the UPR: ATF6, IRE1α, and PERK. Paternal obesity increased GRP78 protein levels in male (but not female) PHF placentae compared to CON placentae at E14.5 (P_Diet_ = 0.0039, P = 0.0022; Fig. 2A) and at E18.5 (P_Sex_ = 0.0247, P_Int_ = 0.0247, P = 0.0407; Fig. 2B). Although *Atf6*, and *Edem1* transcript levels were similar between groups (Fig. 2C-F), *Pdia2* levels were lower in E14.5 PHF female (but not male) placentae compared to CON (P_Diet_ = 0.0139, P_Int_ = 0.0051; P = 0.0038; Fig. 2G), but not at E18.5 (Fig. 2H).

**Figure 2.**
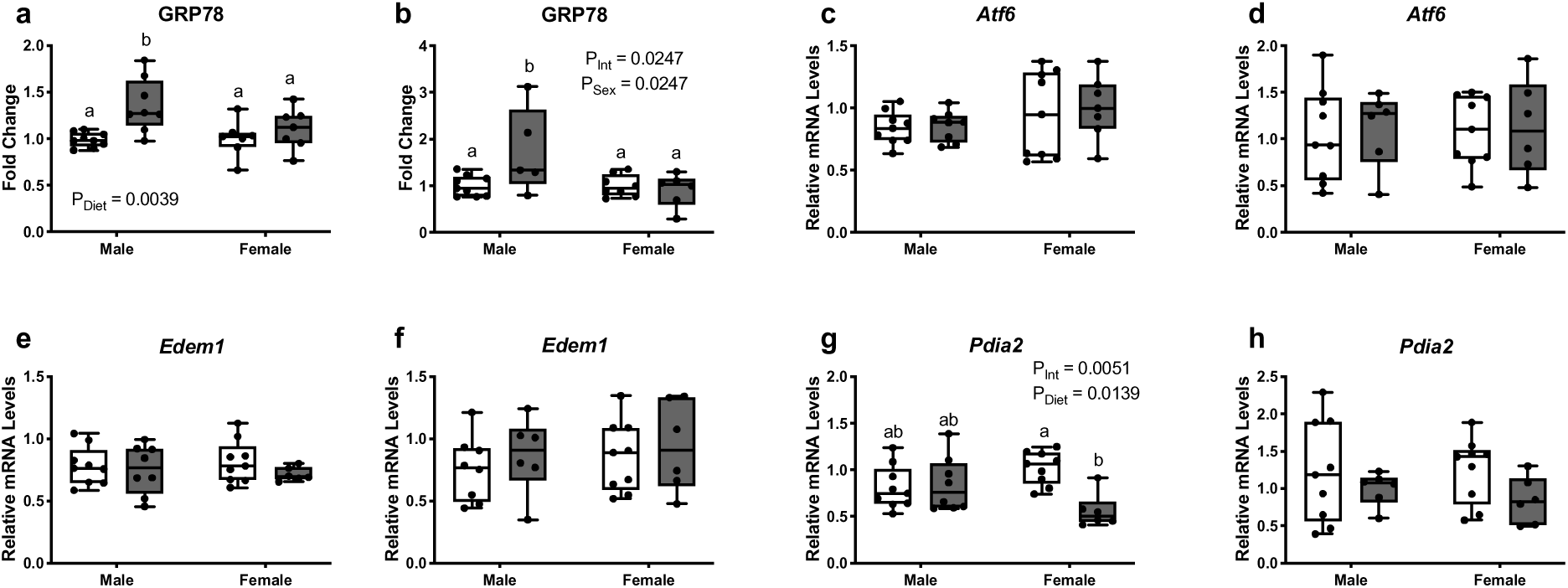
Paternal obesity has sex-specific impacts on endoplasmic reticulum stress in E14.5 and E18.5 placentae. GRP78 protein levels were semi-quantified using Western blotting in CON and PHF E14.5 and E18.5 placentae. All data were normalized to β-actin. Gene expression of *Atf6* pathway and downstream targets were quantified using RT-qPCR in CON and PHF E14.5 and E18.5 placentae. (**a**) GRP78 protein levels in E14.5 and (**b**) E18.5 placentae. (**c**) *Atf6* transcript levels in E14.5 and (**d**) E18.5 placentae. (**e**) *Edem1* transcript levels in E14.5 and (**f**) E18.5 placentae. (**g**) *Pdia* transcript levels in E14.5 and (**h**) E18.5 placentae. CON; n = 7-9, PHF; n = 5-8. Data are presented as box and whisker plots; min to max with a line representing the median. Two-way ANOVA with main effects of paternal diet and placental sex as factors using Bonferroni’s post-hoc for multiple comparison. Box plots with different letters indicate significance *P < 0.05*. CON = Control (open box plots); PHF = Paternal high fat diet-induced obesity (grey box plots).

The IRE1 arm of the UPR is ubiquitously expressed and elicits an adaptive response to ER stress, working to increase ER capacity and decrease ER load.^33^ Paternal obesity increased levels of phosphorylated IRE1α (P_Int_ = 0.0488; Fig. 3A) at E14.5 in female placentae, but were similar between groups at E18.5 (Fig. 3B). Despite this, levels of the downstream transcriptional target *Xbp1*, were similar between groups, including spliced (*Xbp1s*; Fig. 3C,D), total (*Xbp1t;* Fig. 3E,F), and the ratio of *Xbp1s* to *Xbp1t* (*Xbp1s:Xbp1t*; Fig. 3G,H) at E14.5 and E18.5. To understand whether ER stress had downstream functional changes in the placenta, we also investigated whether paternal obesity-induced ER stress activated proinflammatory signaling pathways that involve nuclear factor-κB (NF-κB). At E14.5 and E18.5, NF-κB activity and transcript levels of inflammatory cytokines regulated by NF-κB, including *Traf6*, *Tnf* and *Il1*, were similar between groups (Supplemental Table S5). *Il6* transcript levels however were moderately higher in PHF placental samples compared to CON (Supplemental Table S5). The PERK arm of the UPR inhibits ribosomal function and globally diminishes protein production to accommodate the protein stress adaptation.^33^ Paternal obesity increased the ratio of phosphorylated (phospho-) PERK to total PERK protein levels in E14.5 male (but not female) PHF placentae (P_Int_ = 0.0018, P_Sex_ = 0.0010; P = 0.0356; Fig. 3I) compared to CON. The ratio of phospho-PERK to total PERK protein levels was also increased in PHF female placentae at E18.5 (P_Diet_ = 0.0031, P 0.0277; Fig. 3J). Downstream of PERK activity, the phosphorylation of eukaryotic initiation factor 2A (eIF2α) promotes apoptosis, and the transcriptional activation of pro-apoptotic *Atf4* and *Ddit3* (the gene encoding CHOP) induce proteins involved in amino acid transport, autophagy, folding chaperones, and redox regulatory proteins in addition to pro-apoptotic molecules.^33^ The ratio of phospho-eIF2α to total eIF2α protein levels and transcript levels of *Atf4* and *Ddit3* were unchanged at both time points (Supplemental Table S6). Pro-apoptotic *Bax* transcript levels were unchanged in PHF placentae at E14.5 and E18.5 (Fig. 3K,L); however, anti-apoptotic *Bcl2* transcript levels were increased in female (but not male) PHF placentae at E18.5 (P_Diet_ = 0.0253, P = 0.0312; Fig. 3N) but not E14.5 (Fig. 3M). The ratio of *Bax* to *Bcl2* transcripts levels was unchanged at both time points (Fig. 3O,P). The ratio of cleaved caspase 3 to total caspase protein levels was also unchanged at E14.5 and E18.5 (data not shown). Collectively, these data suggest that paternal obesity does not significantly impact placental apoptosis through the PERK branch of the UPR at E14.5 or E18.5.

**Figure 3.**
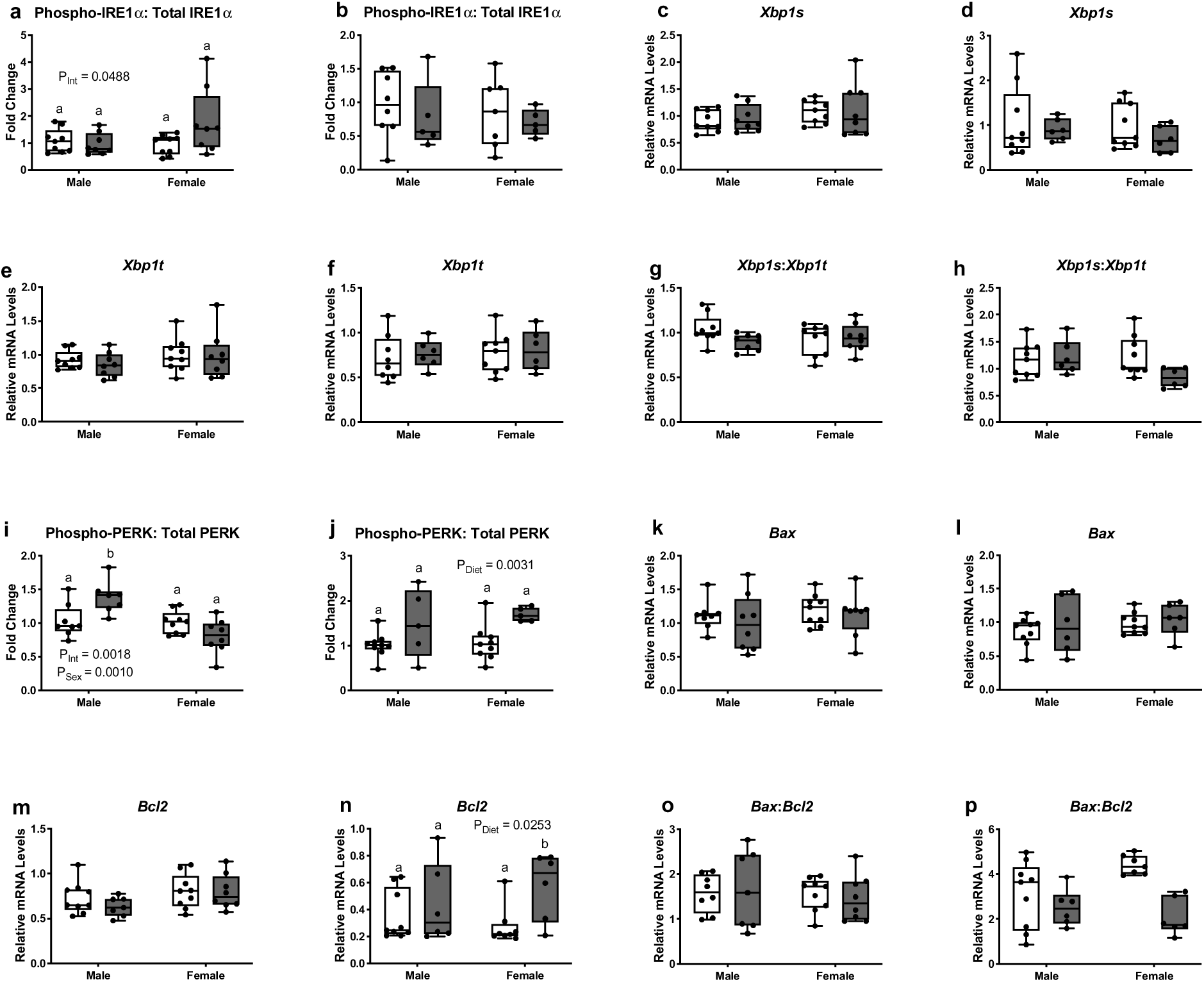
Paternal obesity alters placental endoplasmic reticulum stress branches IRE1α and PERK in a sex-specific manner without changing downstream targets. IRE1α and PERK protein levels were semi-quantified using Western blotting in CON and PHF E14.5 and E18.5 placentae. All data was normalized to β-actin. Gene expression of IRE1α downstream targets and apoptosis-related genes were quantified using RT-qPCR in E14.5 and E18.5 CON and PHF placentae. (**a**) Protein levels of phospho-IRE1α to total IRE1α in E14.5 and (**b**) E18.5 placentae. (**c**) Transcript levels of spliced *Xbp1* (*Xbp1s*) in E14.5 and (**d**) E18.5 placentae. (**e**) Transcript levels of total *Xbp1* (*Xbp1t*) in E14.5 and (**f**) E18.5 placentae. (**g**) Transcript levels of s*Xbp1* to t*Xbp1* in E14.5 and (**h**) E18.5 placentae. (**i**) Protein levels of phospho-PERK to total PERK in E14.5 and (**j**) E18.5 placentae. (**k**) Transcript levels of *Bax* in E14.5 and (**l**) E18.5 placentae. (**m**) Transcript levels of *Bcl2* in E14.5 and (**n**) E18.5 placentae. (**o**) Ratio of transcript levels of *Bax:Bcl2* in E14.5 and (**p**) E18.5 placentae. CON; n = 7-9, PHF; n = 5-8. Data are presented as box and whisker plots; min to max with a line representing the median. Two-way ANOVA with main effects of paternal diet and placental sex as factors using Bonferroni’s post-hoc for multiple comparison. Box plots with different letters indicate significance *P* < 0.05. CON = Control (open box plots); PHF = Paternal high fat diet-induced obesity (grey box plots).

### Paternal obesity alters placental IGF2 and IRS transcripts at mid-gestation

Insulin-like growth factors (IGFs) and their binding to the insulin receptor, engages insulin signaling and regulates placental growth.^34^ Insulin-like growth factor 2 (*Igf2*) is paternally imprinted and in human work, paternal obesity is associated with newborn hypomethylation of insulin-like growth factor 2 (IGF2) protein and ER stress can alter this IGF2 bioactivity.^35^ Therefore we investigated whether paternal obesity-induced ER stress impacted placental IGF in our model. Transcript levels of *Igf2* were elevated in PHF placentae at E14.5 (P_Diet_ = 0.0002, P = 0.0009; Fig. 4A), particularly in placentae associated with female fetuses. This increase however, did not persist to E18.5 (Fig. 4B). IGF2 binds to IGF-1R, IGF-2R and insulin receptors (IR)^36^ and targets insulin receptor substrate 1 (IRS1) and IRS2, to promote placental growth and development through cell proliferation, survival, and mitogenesis.^37^ Consistent with increased *Igf2* levels, *Irs1* transcripts were elevated in PHF placentae compared to CON (P_Diet_ = 0.0483, P_Sex_ = 0.0041; Fig. 4C) at E14.5 but were similar between groups at E18.5 (Fig. 4D). Transcript levels of *Irs2* were similar between groups at E14.5 (Fig. 4E), but at E18.5 an interaction was noted where transcript levels were lower in PHF male placentae and higher in PHF female placentae (P_Int_ = 0.0068; Fig. 4F).

**Figure 4.**
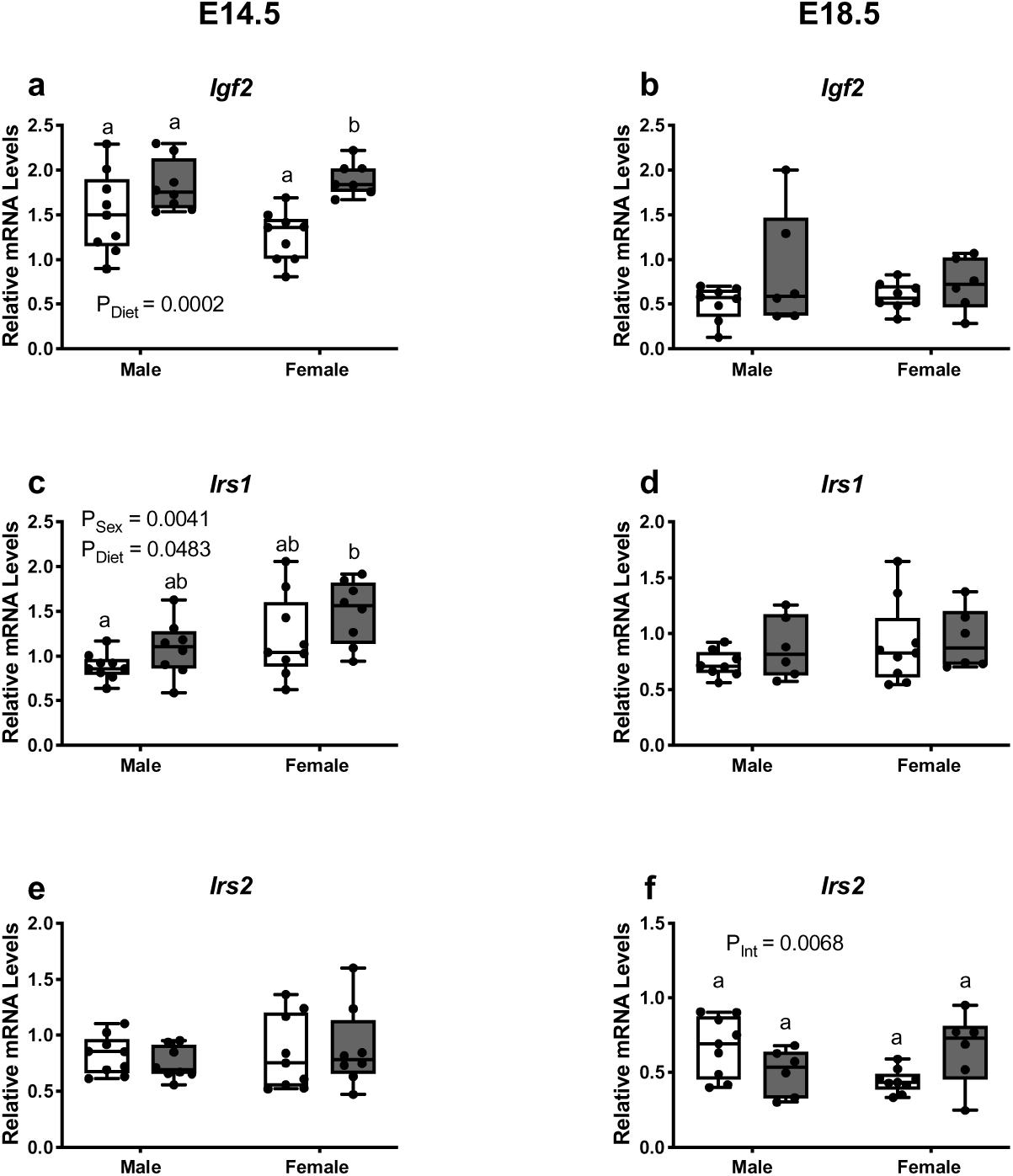
Paternal obesity alters placental *Igf2* transcripts and its downstream signaling factors in a sex-specific manner. Gene expression of key regulators of IGF2-IRS signaling were quantified using RT-qPCR in CON and PHF E14.5 and E18.5 placentae. (**a**) Transcript levels of *Igf2* in E14.5 and (**b**) E18.5 placentae. (**c**) Transcript levels of *Irs1* in E14.5 and (**d**) E18.5 placentae. (**e**) Transcript levels of *Irs2* in E14.5 and (**f**) E18.5 placentae. CON; n = 8-9, PHF; n = 6-8. Data are presented as box and whisker plots; min to max with a line representing the median. Two-way ANOVA with main effects of paternal diet and placental sex as factors using Bonferroni’s post-hoc for multiple comparison. Box plots with different letters indicate significance *P* < 0.05. CON = Control (open box plots); PHF = Paternal high fat diet-induced obesity (grey box plots).

### Despite changes to placental vasculature and *Igf2* transcripts, paternal obesity does not impact placental endocrine-mediated maternal glucose metabolism

The placenta is responsible for the secretion of a number of endocrine factors, including placental lactogen, which facilitates maternal metabolic adaptations to pregnancy.^38^ Since we observed significant changes in placental development in pregnancies sired by PHF, we determined whether placental-mediated changes in maternal glucose tolerance were impacted through changes to key cell types that produce placental lactogen, or whether placental lactogen gene transcripts were altered. No studies to date have investigated whether paternal diet/obesity can affect maternal metabolic adaptations during pregnancy, and thus we also investigated glucose tolerance at E14.5 and E18.5 in females mated with CON and PHF males. We observed that at both time points in pregnancy, maternal glucose tolerance is unaltered (Fig. 5A-D). This *in vivo* observation is supported by our histological analyses of parietal trophoblast giant cells numbers (Fig. 5E-G). Similarly, maternal serum insulin and HOMA-IR was also similar between groups (data not shown). Transcript levels of placental lactogen I coding-gene (*Csh1)* were unaltered at E14.5 (Fig. 5H), although at E18.5 transcript levels were lower in female placentae compared to male placentae (P_Sex_ = 0.0021; P = 0.0140; Fig. 5I). Expression levels of *Csh2* were similar between groups at E14.5 (Fig. 5J) and E18.5 (Fig. 5K). Our findings suggest that maternal glucose tolerance and insulin sensitivity, as well as the capacity to produce placental lactogen at mid-gestation and term gestation was unaffected by obesity in the father.

**Figure 5.**
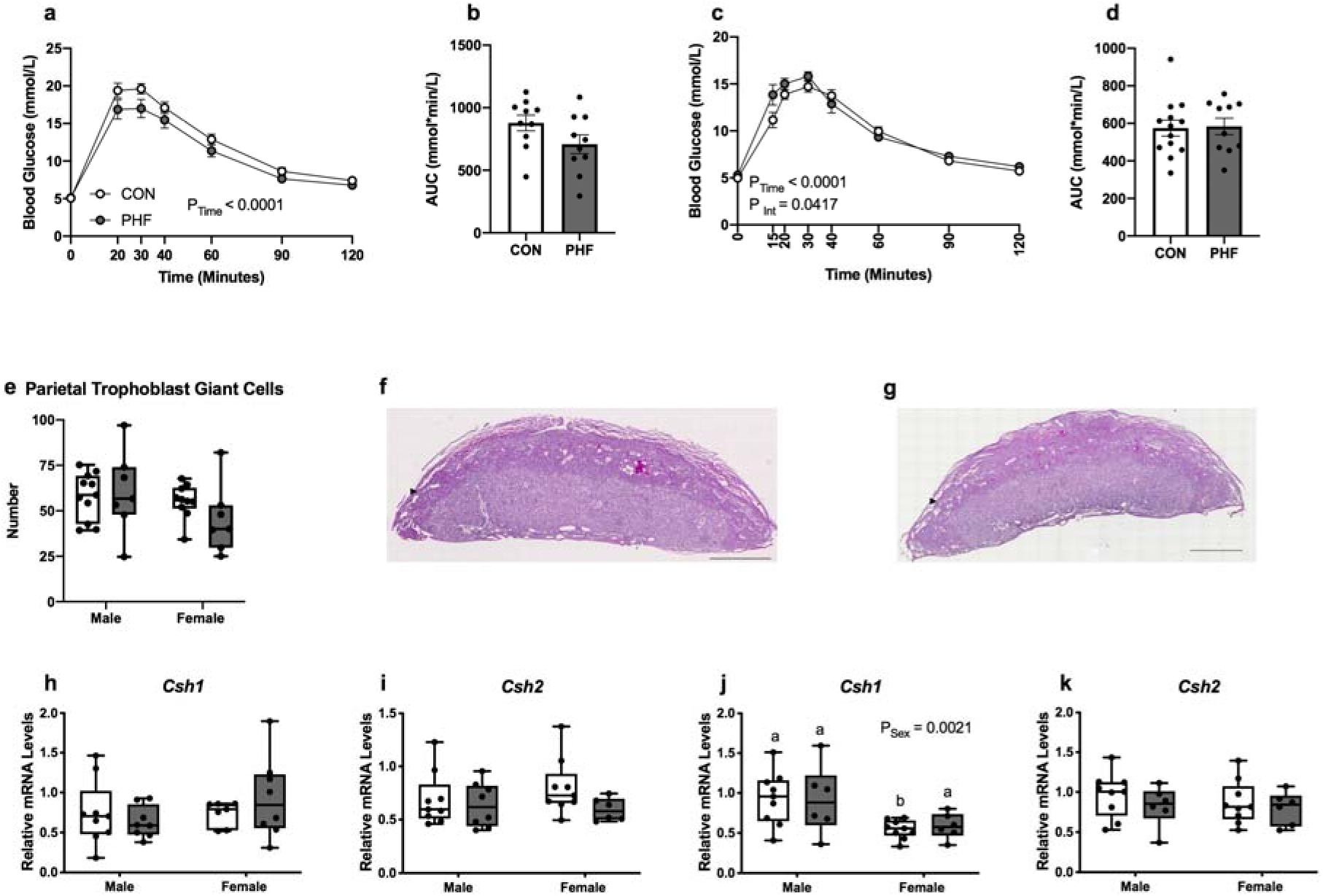
Paternal obesity has no effect on maternal glucose tolerance or placental lactogen at mid-gestation and term gestation. Female mice mated with either CON or PHF males were subjected to a standard glucose tolerance test to assess glucose tolerance at mid-gestation (E14.5, 1.5g/kg *i.p.*) or term gestation (E18.5, 2g/kg gavage). (**a**) Maternal glucose tolerance, and (**b**) glucose area under the curve (AUC) was similar between groups at E14.5. (**c**) Maternal glucose tolerance, and (**d**) glucose AUC was similar between groups at E18.5. (**e**) Number of parietal trophoblast giant cells in CON and PHF placentae. (**f**) Representative photos of E14.5 CON and (**g**) PHF placental samples stained for glycogen using PAS. Arrowheads show placental parietal trophoblast giant cells stained with Periodic Acid Schiff (pink). (**h**) *Csh1* transcript levels at E14.5 and at (**i**) E18.5. (**j**) *Csh2* transcript levels at E14.5 and at (**k**) E18.5. Data are presented as mean ± SEM for glucose tolerance, and as box and whisker plots; min to max with a line representing the median for transcript levels. Glucose tolerance was measured using two-way repeated measures ANOVA or mixed-effects model with main effects of paternal diet and time as factors using Bonferroni’s post-hoc for multiple comparison; or by unpaired Student’s t-test. Placental *Csh* transcripts were measured using two-way ANOVA with main effects of paternal diet and placental sex as factors using Bonferroni’s post-hoc for multiple comparison. Box plots with different letters indicate significance *P* < 0.05. CON = Control (open circles and box plots); PHF = Paternal high fat diet-induced obesity (grey circles and box plots).

### Paternal obesity alters placental amino acid system A nutrient transporter expression

Appropriate placental vessel development is critical for nutrient transport. Since we observed changes that were indicative of placental hypoxia and altered vessel development, we next investigated levels of key placental nutrient transporters. Transcript levels of *Slc38a2*, which encodes sodium-coupled neutral amino acid protein 2 (SNAT2), were similar between groups at E14.5 (Supplemental Table S7), but were higher in female PHF placentae compared to CON at E18.5 (P_Int_ = 0.0282, P_Diet_ = 0.0837; P = 0.0493; Supplemental Table S7). In contrast, transcript levels of genes that encode glucose transporter 1 (GLUT1, *Slc2a1*) and GLUT3 (*Slc2a3*) were similar between groups (Supplemental Table S7), as were transcript levels of *Fabp4*, which encodes fatty acid binding protein (Supplemental Table S7). These data suggest that paternal obesity may change amino acid transport at term gestation but does not appreciably impact most placental nutrient transporter levels.

### Paternal obesity impairs overall mating efficiency but does not impact fetal growth or pregnancy outcomes

Diet-induced obesity in males is known to decrease sperm count and sperm motility, reduces number of sperm with normal morphology,^39^ increases sperm oxidative stress-induced DNA damage,^40^ and is associated with reduced fertility.^39^ We found that more male-female pairings were necessary to produce a pregnancy in females mated with PHF males compared to females mated with CON males (P = 0.0424; Fig. 6A). We also observed that there were significantly fewer pregnancies generated as a proportion of total copulation plugs in PHF-mated females (P = 0.0006; Fig. 6B). There were also significantly less copulation plugs generated as a proportion of the total number of pairings with PHF males (P = 0.0089; Fig. 6C), and that it took moderately longer for females paired with PHF males to produce a copulation plug (Fig. 6D), although this difference was not statistically significant (p = 0.0856).

**Figure 6.**
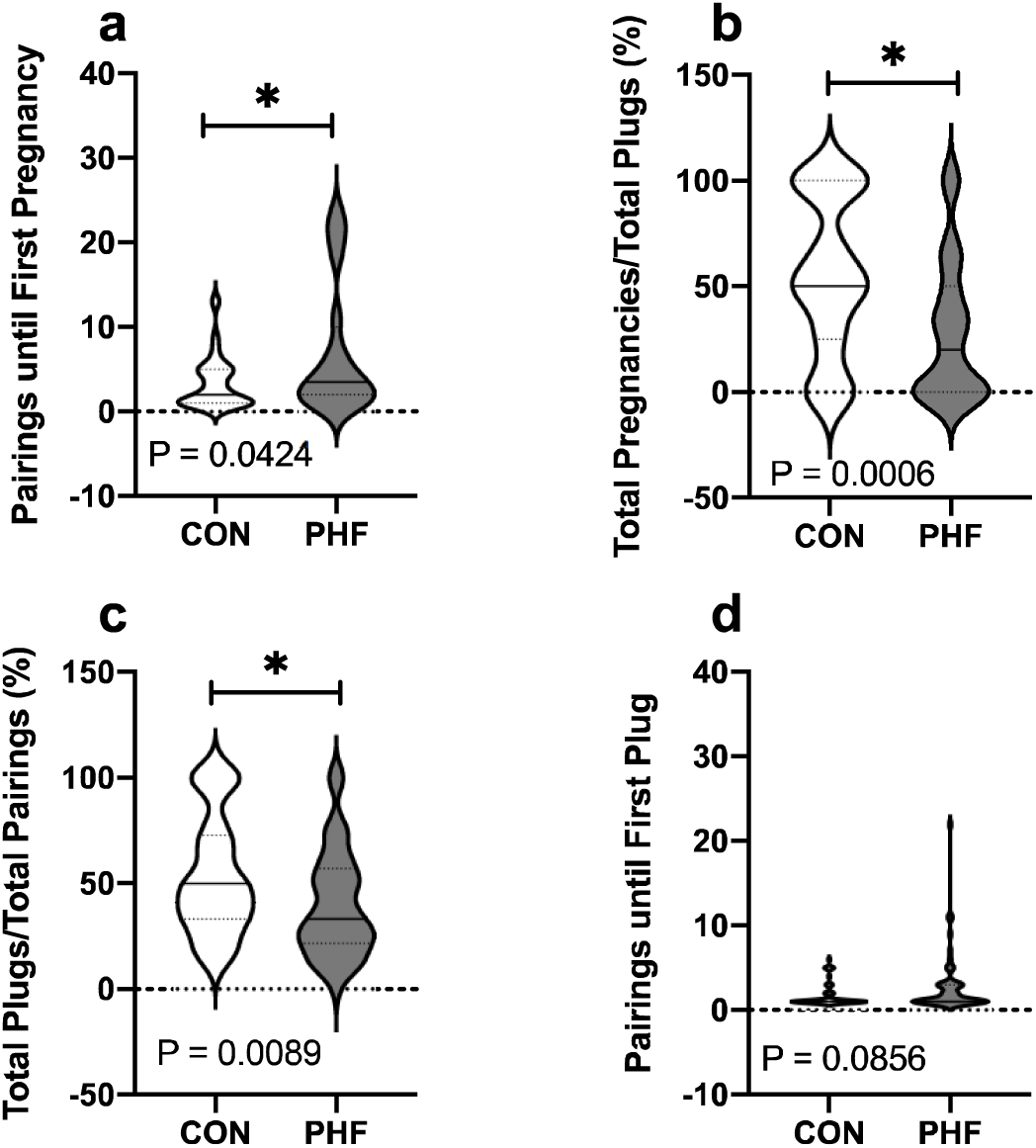
Paternal obesity impairs mating efficiency. CON and PHF males were time-mated with females to generate pregnancies. (**a**) More mating attempts were necessary to produce a pregnancy in females mated with PHF males compared to females mated with CON males. (**b**) There were significantly fewer pregnancies generated as a proportion of total copulation plugs in PHF-mated females. (**c**) Fewer copulation plugs were generated as a proportion of total pairings in PHF-mated females. (**d**) It took moderately longer for females paired with PHF males to produce a copulation plug compared to those paired with a CON male. Data are presented as volcano plots with the center line representing the median. \**P* < 0.05 compared to CON using Student’s t-test or Mann-Whitney U test. CON = Control (open plots); PHF = Paternal high fat diet-induced obesity (grey plots).

Although evidence from rodent models suggests that paternal obesity reduces fertilization success, reduces implantation rate,^10^ and alters fetoplacental development, ^10^ in our study, maternal body mass gained during gestation and fasting blood glucose levels, litter size, number of resorptions per litter, and fetal sex ratio were similar between CON and PHF dams at E14.5 and E18.5 (Supplemental Table S8). CON female fetal body mass was significantly lower compared to CON male fetal body mass (P = 0.0079) at E14.5; however, placental mass was unchanged at E14.5 (Supplemental Table S8). Placental and fetal body mass at E18.5 were also similar between male and female CON and PHF litters (Supplemental Table S8).

### Paternal obesity promotes fetal hepatic ER stress and activation of gluconeogenesis at term gestation

It has been previously shown that paternal obesity is associated with changes in glucose tolerance in adult offspring, particularly in females.^33^ Since fetal hepatic gluconeogenesis contributes to postnatal glucose tolerance, and maternal obesity is associated with changes in fetal gluconeogenic enzyme expression,^5^ we investigated whether paternal obesity similarly impacted fetal hepatic metabolic development. Hepatocyte nuclear factors (HNFs) play an important role in embryonic hepatic development. Hepatocyte nuclear factor 4 alpha (HNF4A) is a key regulator of many genes involved in hepatic function, including apolipoproteins and its targeted deletion in the liver results in steatosis, and severe disruption of gluconeogenesis.^41^ Although transcript levels of the *Hnf4a* were unaltered at E14.5 (Fig. 7A), levels were higher in PHF male (but not female) livers compared to CON at E18.5 (P_Int_ = 0.0229, P_Sex_ = 0.0008, P_Diet_ < 0.0001; Fig. 7B). Transcript levels of the HNF4A co-activator, peroxisome proliferative activated receptor gamma coactivator 1-alpha (*Ppargc1a*), were also similar between groups at E14.5 (Fig. 7C), but increased at E18.5, albeit only in female PHF livers (P_Diet_ = 0.0213, P = 0.0334; Fig. 7D). We next quantified transcript levels of hepatic glucose-6-phosphatase (*G6p*), which hydrolyzes glucose-6-phosphate and frees glucose. Although overall hepatic levels of *G6p* were higher in females compared to males at E14.5 (P_Sex_ = 0.0216; Fig. 7E), PHF did not affect *G6p* transcript levels at E18.5 (Fig. 7F). Levels of the rate-limiting enzyme in gluconeogenesis, phosphoenolpyruvate carboxykinase 1 (*Pck1*), were similar between groups at both time points. These results demonstrate that expression of hepatic gluconeogenic transcriptional activators are increased in PHF fetuses at E18.5, but key downstream gluconeogenic enzymes are unaffected.

**Figure 7.**
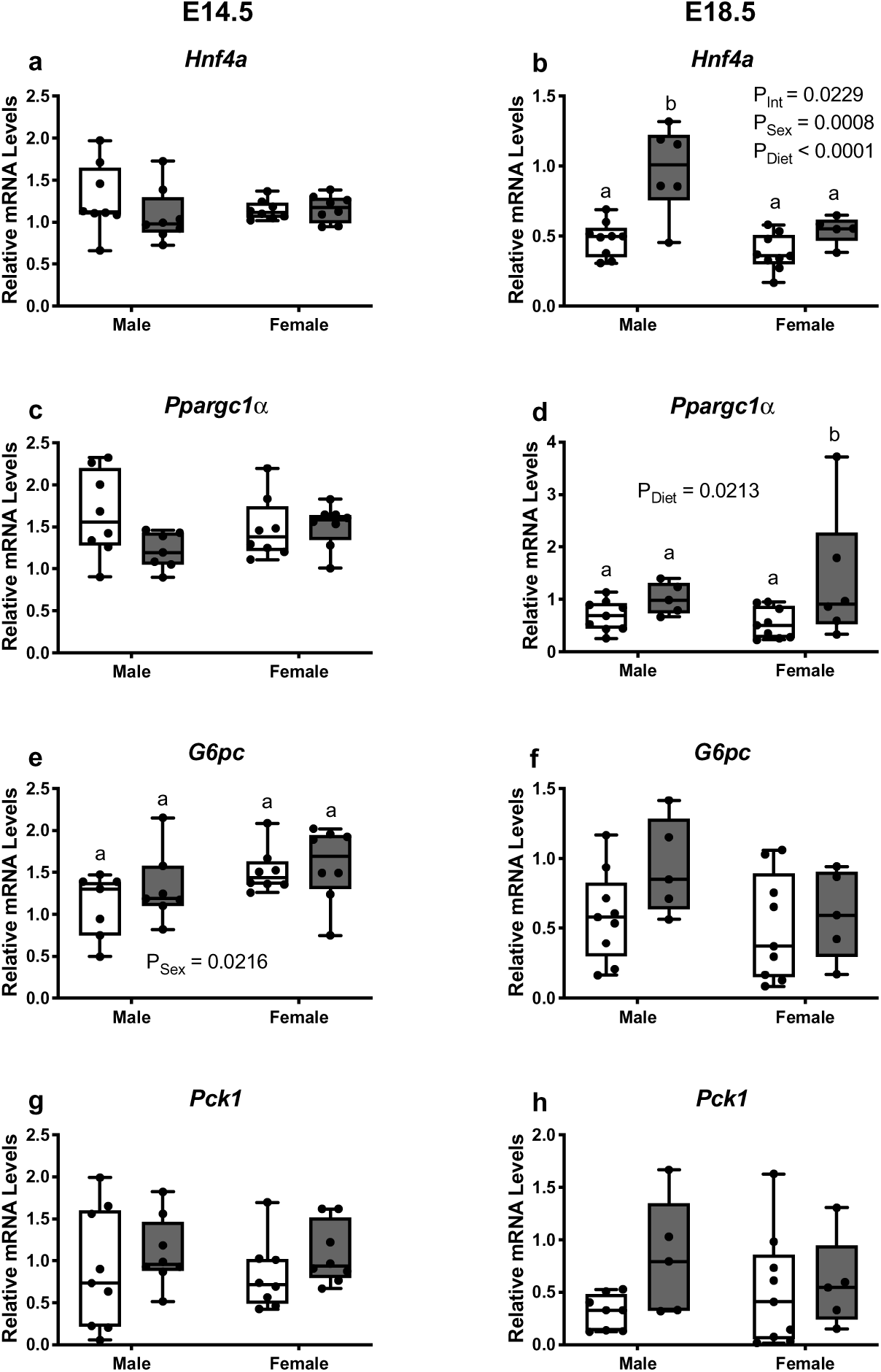
Paternal obesity increased expression of fetal hepatic gluconeogenic transcriptional activators at term gestation. Transcript levels of gluconeogenic transcriptional activators and downstream effectors were quantified using RT-qPCR in CON and PHF E14.5 and E18.5 fetal liver. (**a**) *Hnf4a* transcript levels at E14.5 and (**b**) E18.5 liver. (**c**) *Ppargc1a* transcript levels in E14.5 and (**d**) E18.5 liver. (**e**) *G6pc* transcript levels in E14.5 and (**f**) E18.5 liver. (**g**) *Pck1* transcript levels in E14.5 and (**h**) E18.5 liver. CON; n = 7-9, PHF; n = 5-8. Data are presented as box and whisker plots; min to max with a line representing the median. Two-way ANOVA with main effects of paternal diet and fetal sex as factors using Bonferroni’s post-hoc for multiple comparison. Box plots with different letters indicate significance *P* < 0.05. CON = Control (open box plots); PHF = Paternal high fat diet induced obesity (grey box plots).

### Paternal obesity activates the UPR in the fetal liver

Maternal obesity has been associated with ER stress in offspring.^42^ We investigated whether paternal obesity alters hepatic metabolism through ER stress-induced effectors. Transcript levels of *Hspa5*, the gene that encodes for ER stress chaperone GRP78, were increased in female PHF livers compared to CON at E14.5 (P_Diet_ = 0.0013, P = 0.0202; Fig. 8A), but at term gestation, GRP78 protein levels were unchanged (data not shown). Hepatic expression of *Eif2ak3*, the gene that encodes PERK, was increased in PHF fetuses at E14.5 (CON male; 1.162 ± 0.189, CON female; 1.118 ± 0.219, PHF male; 1.930 ± 0.215, PHF female; 1.533 ± 0.366; P_Diet_ = 0.0333). But, unfortunately, due to lack of sufficient sample, we were unable to measure transcript levels of *Eif2ak3* at E18.5. However, Western blot analyses showed that protein levels of fetal hepatic phospo-PERK at E18.5 were increased male PHF fetuses but not females (P_Diet_ = 0.0018; P = 0.0127; Fig. 8B). Activation of the PERK arm of the UPR was associated with increased transcript levels of pro-apoptotic factor, *Ddit3*, the CHOP-encoding gene, in fetal livers at both E14.5 (P_Diet_ = 0.0075, P = 0.0234; Fig. 8C) and E18.5 (P_Diet_ = 0.0126, P_Sex_ = 0.0023; Fig. 8D), suggesting that the process of apoptosis is elevated in PHF fetal livers. Consistent with this, protein levels of cleaved caspase 3 (CON male; 1.051 ± 0.119, CON female; 1.003 ± 0.090, PHF male; 1.811 ± 0.643, PHF female; 1.595 ± 0.402; P_Diet_ = 0.0347) were also increased in PHF livers in E18.5, although transcript levels of *Bax* were unchanged at E14.5 (Fig. 8E) and decreased in female liver (P_Sex_ = 0.0378; Fig. 8F) at E18.5. Expression of *Bcl2* was increased in PHF male livers compared to CON male livers (P_Diet_ = 0.0140, P = 0.0120; Fig. 8G) at E14.5, but these changes did not persist until E18.5 (Fig. 8H). The ratio of *Bax* to *Bcl2* transcript levels were similar between groups (data not shown).

**Figure 8.**
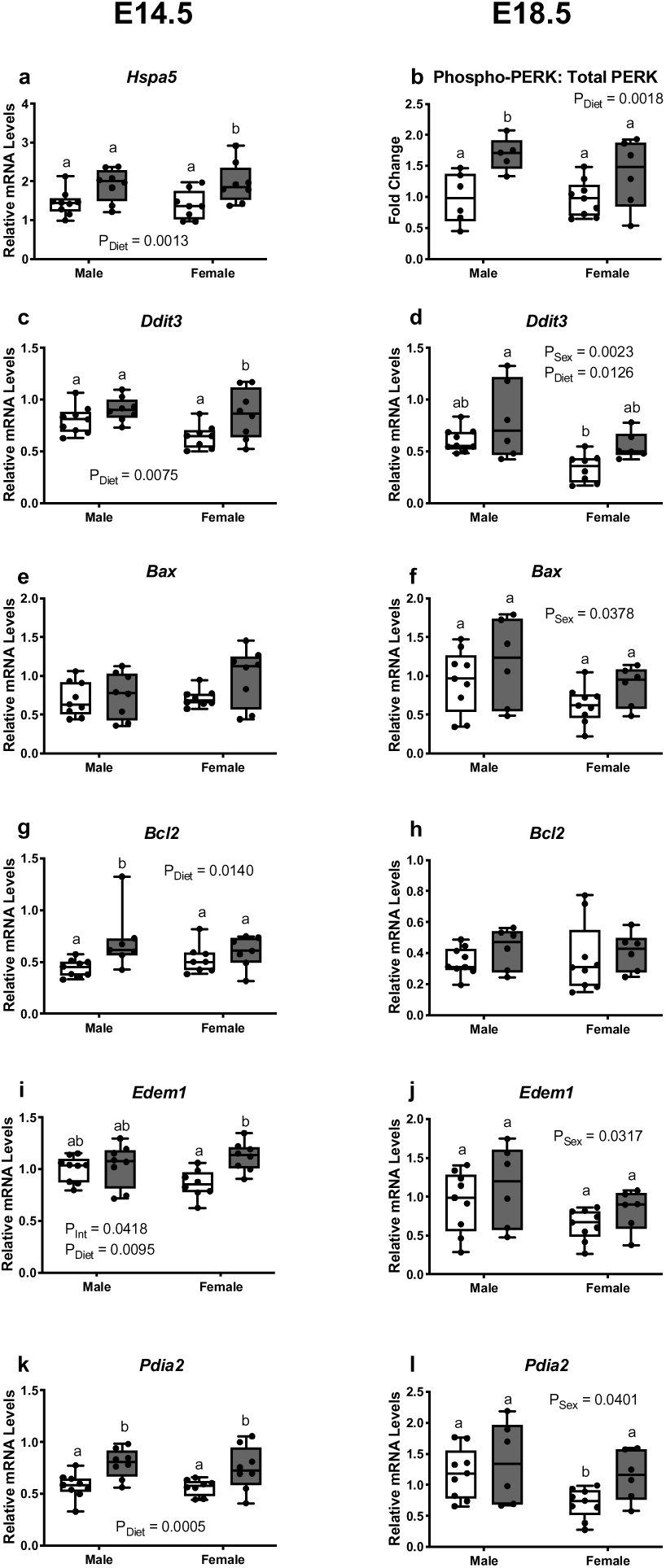
Paternal obesity activates the unfolded protein response in fetal liver. Transcript levels of *Hspa5* and PERK and apoptosis-related genes were quantified using RT-qPCR in CON and PHF E14.5 and/or E18.5 liver. Ratio of protein levels of phospho-PERK:total PERK and cleaved caspase were semi-quantified using Western blotting in CON and PHF E18.5 liver. All data was normalized to β-actin. (**a**) *Hspa5* transcript levels in E14.5 liver. (**b**) Phospho-PERK: Total PERK protein levels in E18.5 liver. (**c**) *Ddit3* transcript levels in E14.5 and (**d**) E18.5 liver. (**e**) *Bax* transcript levels in E14.5 and (**f**) E18.5 liver. (**g**) *Bcl2* transcript levels in E14.5 and (**h**) E18.5 liver. (**i**) *Edem1* transcript levels in E14.5 and (**j**) E18.5 liver. (**k**) *Pdia2* transcript levels in E14.5 and (**l**) E18.5 liver. CON; n = 6-9, PHF; n = 5-8. Data are presented as box and whisker plots min to max; with a line representing the median. Two-way ANOVA with main effects of paternal diet and fetal sex as factors using Bonferroni’s post-hoc for multiple comparison. Boxes with different letters indicate significance *P* < 0.05. CON = Control (open box plots); PHF = Paternal high fat diet-induced obesity (grey box plots).

The native ER protein, ATF6, controls one of the three branches of the UPR and has been shown to be assisted by EDEM1. ATF6 is selectively translated by the phosphorylation of eIF2α and participates on a feedback loop by inducing the expression of the eIF2α phosphatase, GADD34.^43^ While transcript levels of *Ppp1r15a*, the gene that encodes for GADD34, were not changed in PHF liver at E14.5 or E18.5 (data not shown), transcript levels of *Edem1* were increased in female PHF liver compared to female CON liver at E14.5 (P_Int_ = 0.0418, P_Diet_ = 0.0095, P = 0.0112; Fig. 8I) but not at E18.5. In general, *Edem1* transcripts were lower in female compared to male livers at E18.5 (P_Sex_ = 0.0317; Fig. 8J). In addition to the main branches of the UPR, there are a number of genes that have crucial functions in protein folding that are altered with ER stress. PDIA2 is responsible for protein folding and thiol-disulfide exchanges and is mediated by transcription factor AFT6.^44^ Transcript levels of *Pdia* were increased in PHF (P_Diet_ = 0.0005) male (P = 0.0094; Fig. 8K) and female (P = 0.0366; Fig. 8K) livers at E14.5, but at E18.5, *Pdia* expression levels were decreased in female CON liver compared to male CON liver (P_Sex_ = 0.0401, P = 0.0477; Fig. 8L).

Finally, we investigated whether the IRE1 branch of the UPR was altered in fetal liver due to paternal obesity. Neither IRE1α transcript nor protein levels were altered in CON and PHF E14.5 and E18.5 (data not shown) liver; the ratio of *Xbp1s*:*Xbp1t* was similar between groups. Thus, although PERK and ATF6 arms of the UPR are activated in fetal liver in response to paternal obesity, the IRE1 branch is not.

### Paternal obesity is associated with changes in whole body energetics and results in glucose intolerance in offspring

Paternal obesity has been shown to significantly impact offspring metabolism.^7^ In our study, CON and PHF male and female offspring (Supplemental Fig. S3A,B) gained similar amounts of weight and ate a similar number of calories daily (Supplemental Fig. S3C,D). Offspring gonadal fat mass was affected by both offspring sex and paternal diet (P_Sex_ < 0.0001, P_Int_ = 0.0153; Supplemental Fig. S3E); PHF male offspring had lower fat mass, and PHF females had higher fat mass. Females in general had lower gonadal fat mass compared to males (Supplemental Fig. S3E). Although paternal obesity did not impact liver weight, females had lighter livers compared to male offspring (P_Sex_ < 0.0001, P = 0.0007; Supplemental Fig. S3F).

To our knowledge, no studies exist that have investigated whether paternal obesity alters whole body energy metabolism in offspring. We used a sophisticated metabolic assessment system that uses indirect calorimetry^45^ to calculate whether paternal obesity impacted the type and rate of substrate utilization by the offspring, while also measuring 48hr activity and food consumption over both the light and dark cycles. Food consumption over a 48hr period, was higher in PHF female and male offspring (P_Diet_ = 0.011; Fig. 9A) and was especially higher during the dark compared to light cycle of the day. Heat production, a marker of overall energy consumption, trended to be lower in PHF offspring (P_Diet_ = 0.055; Fig. 9B) and was greater so in female offspring. Total activity over 48hrs was lower in male offspring (P_Sex_ = 0.041; Fig. 9C) compared to females overall and tended to be lower in PHF females compared to CON – particularly during the light cycle. Oxygen consumption (VO_2_ = O_2_ inhaled per unit time) was lower in male PHF offspring (P_Sex_ < 0.0001; Fig. 9D), as was CO_2_ production (VCO_2_ = CO_2_ exhaled from the body per unit time; P_Sex_ = 0.004; Fig. 9E), and tended to be lower in females-again especially during the light cycle. Using VO_2_ and VCO_2_ we calculated the RER to estimate the respiratory quotient (RQ), an indicator of which fuel (e.g. carbohydrate or fat) is being metabolized^45^ over a 48hr period. An RER value near 0.7 suggests that fat is the predominant fuel source, a RER value near 1.0 suggests that carbohydrate is the predominant fuel source, and a RER value between 0.7 and 1.0 suggests a combination of fat and carbohydrate is being used. Although RER trended to be lower in PHF offspring during the light cycle, but this difference was not statistically significant (Fig 9F). Lipid oxidation was lower in male offspring (P_Sex_ = 0.012; Fig. 9G), and water consumption was similar between CON and PHF groups (data not shown). Thus, our data suggest that basal metabolic rate appears moderately lower in PHF offspring, and that impacts of paternal obesity on whole body energy metabolism in offspring is also be sex-specific.

**Figure 9.**
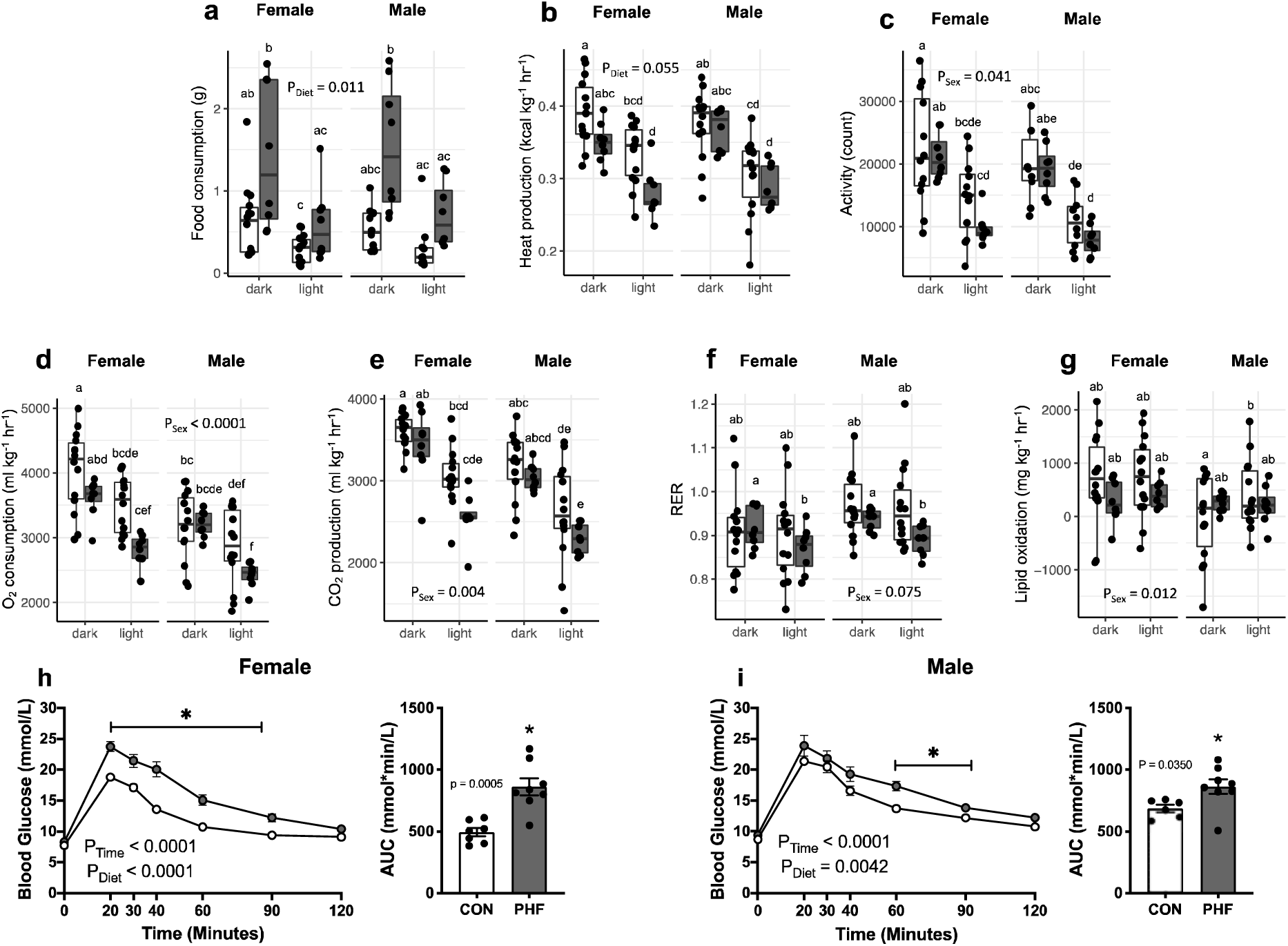
Paternal obesity results in altered whole body energy metabolism and glucose intolerance in offspring. Young adult CON and PHF male and female offspring were subjected to *in vivo* metabolic assessment at P53 and a glucose tolerance test (2g/kg, *i.p.*) at P60. (**a**) Adult CON and PHF male and female offspring food consumption, (**b**) heat production, (**c**) total horizontal motor activity, (**d**) oxygen consumption, (**e**) carbon dioxide production, (**f**) respiratory exchange ratio, and (**g**) lipid oxidation. Metabolic assessments data are presented as box and whisker plots; min to max with a line representing the median. Glucose tolerance test and AUC are presented as mean ± SEM. Two-way repeated measures ANOVA with main effects of paternal diet and time as factors using Bonferroni’s post-hoc for multiple comparison or Student’s t-test. Metabolic assessments presented as box plots with different letters denote *P* < 0.05 in a linear mixed-effects model with main effects and multiple comparisons determined by Satterthwaite’s method of approximation. CON = Control (open circles and box plots); PHF = Paternal high fat diet-induced obesity (grey circles and box plots).

Previous studies have shown that paternal diet impairs offspring glucose tolerance and may be associated with impaired insulin secretion in offspring.^7^ In our study, paternal obesity increased fasting blood glucose levels in male and female offspring at P60 (CON male; 8.7 ± 0.2, CON female; 7.6 ± 0.1, PHF male; 9.5 ± 0.2, PHF female; 8.1 ± 0.3, P_Diet_ = 0.017, P_Sex_ < 0.0001), although multiple comparisons between CON and PHF males and females did not reach statistical significance (data not shown). We subjected young adult (P60) female and male offspring sired by either CON or PHF males to a glucose challenge via a GTT. Paternal obesity impaired glucose tolerance in both female (P_Diet_ < 0.0001; Fig. 9H) and male (P_Diet_ = 0.0042; Fig. 9I) offspring at P60. Thus, paternal obesity is associated with changes in both whole body metabolism and glucose intolerance in young adult offspring.

## DISCUSSION

The first experimental data showing that paternal diet/obesity results in glucose intolerant offspring were published over a decade ago.^7^ Yet despite this, we still understand very little of the mechanisms underpinning this relationship. To understand the early life regulatory factors that contribute to offspring metabolic compromise, we investigated the effects of paternal obesity on maternal and fetal glucose metabolism, and placental vascular development. We demonstrate that not only is paternal obesity is associated with reduced mating efficiency in sires, it induces placental hypoxia and ER stress, and altered placental angiogenesis. We show that placentae associated with a female fetus are especially vulnerable; changes to transcription factors that regulate placental vascular remodeling were associated with activation of the PERK and IRE1α arms of the UPR, and with increased *Igf2* and changes in the amino acid transporter SNAT2. We also show that paternal obesity has significant impacts on fetal metabolism, inducing hepatic ER stress through the PERK and ATF6 arms of the UPR in both male and female fetuses. These changes to the fetus and placenta, however, were not accompanied by changes in fetal body mass, sex ratio, or litter size. To date, no one has investigated whether paternal obesity may have indirect impacts on offspring through changes to placental development and downstream changes in maternal metabolic function. We now show that despite significant changes to the placenta, paternal obesity does not affect maternal glucose metabolism or the number of trophoblast giant cells that produce placental lactogen. Despite this, placental and fetal hepatic impairments still result in glucose intolerance and impair whole body energetics in offspring at a relatively young age. We suggest that together, intrauterine alterations in placental and fetal hepatic function underpin metabolic dysfunction in male and female offspring and changes in offspring metabolic rate, changes that appear fueled by ER stress *in utero*.

### Paternal obesity impacts placental vascular development and results in ER stress

Placental hypoxia is often associated with pregnancy complications, including preeclampsia,^32^ with poor fetal and offspring outcomes.^46^ Indeed, experimental studies have shown that maternal obesity is associated with impaired placental vascularization, increased ER stress, and inflammation.^4, 5^ To our knowledge, our study is the first to show that paternal obesity (independent of maternal obesity) can induce changes in placental blood vessel development and hypoxia. Our observations that placentae of fetuses sired from obese fathers had elevated HIF-1α protein levels and pro-angiogeneic factors, particularly in females, are consistent with human observations in preeclamptic placentae,^46^ and consistent with the hypothesis that hypoxia can activate a sequence of events starting with upregulation of HIF-1α production. ^47^ Recently, it has been suggested that a decrease in the transcription factor HMOX1 may impair the HIF-1α response to hypoxia-reoxygenation in placentae, also impairing downstream placental vascular formation.^48^ Whether this transcriptional relationship governs the outcomes that we see here is unclear, although our data do suggest that the link between HMOX1 and HIF-1α and their impacts on VEGF likely plays a key role in paternal obesity-induced impaired placental vascular development.

We investigated whether this altered relationship between VEGF, HIF-1α and hypoxia could be mediated by placental ER stress. Hypoxia and ER stress are known to independently promote VEGF expression,^49^ where ER stress enhances the phosphorylation of HIF-1α and potentiates HIF-1α-induced VEGF expression.^50^ We show paternal obesity is associated with placental activation of the PERK arm of the UPR and increased levels of GRP78 in male placentae only. We believe that ER stress could be driving compensatory angiogenic adaptations these PHF placentae. We observe an increase in phospho-PERK protein levels at term in PHF placentae, which was associated with increased anti-apoptotic marker *Bcl2*. Anti-apoptotic BCL2 activity has been linked to maintenance of angiogenesis with HIF-1α and VEGF via PERK branch of the UPR.^51^ A similar relationship may exist in placentae derived from obese fathers.

IRE1α appears necessary for placental labyrinth VEGF expression.^31^ We observe an increase in IRE1α in female PHF placentae at mid-gestation, but whether this increase directly participates in VEGF induction is unknown. Assuming conventional UPR signaling pathways are operating in the placentae, and since we show increased levels of phospho-PERK and phospho-IRE1α, we would expect increased levels of pro-inflammatory cytokines induced by IRE1α,^52^ and hypoxia.^53^ Although we see a moderate increase in *Il6* transcripts at E18.5 in PHF placentae, there was overall little evidence of inflammation in placentae from obese fathers. Although the low levels of ER stress observed may not be robust enough to drive profound inflammation, we do however observe an imbalance of pro-apoptotic and anti-apoptotic signaling in PHF placentae. This is supported by decreased transcript levels of the gene encoding MMP14, evident in PHF placentae. MMPs break down the extracellular matrix between endothelial cells to promote blood vessel widening during proper vascular development^54^ and are mediated by hypoxia;^55^ and indeed, lower transcript levels have been shown to be associated with preeclampsia.^56^ It appears likely that the signaling pathways that drive blood vessel development in the placenta are altered with paternal obesity, and we suggest that the upstream initiator may be placental ER stress.

A primary function of the placenta is to transport nutrients from the mother to the fetus, and many studies show that disruptions in placental development impair nutrient transport leading to intrauterine growth restriction.^17^ Although we do not show changes in many nutrient transporter with paternal obesity, we observe higher levels of Slc3*8a2* (SNAT2) in female PHF placentae at E18.5. SNAT2 transports neutral amino acids, including alanine and glutamine,^57^ which are particularly important as they are glucogenic amino acids and can be converted to glucose when needed.^58^ Although the upstream drivers of this increase are not established, SNAT2 is known to be stimulated by both osmoregulatory and cellular stress regulators.^59^ Osmotic stress initiates processes including cell shrinkage, oxidative stress, and DNA damage^60^ and in placental trophoblast, *in vitro* hypertonicity increases SNAT2 expression.^61^ Furthermore, SNAT2 is the most frequently induced amino acid transporter by hypoxia in breast cancer, and appears regulated by hypoxia both *in vitro* and *in vivo* in cancer models.^62^ Together, these data suggest that a relationship between cellular stress, SNAT2, and angiogenic regulators could exist in the placentae in our model of paternal obesity.

Placental growth and development are regulated by insulin-like growth factors (IGFs) via both endocrine and substrate regulatory pathways.^34^ We show increased *Igf2* transcript levels in PHF placentae, consistent with increased levels of *Irs1*, but not *Irs2*; this suggests that in our model, IGF2 may be signaling through IGF1R pathways. Indeed, IGFs have both proliferative and anti-apoptotic effects on trophoblast through activating IGF1R and triggering the MAPK and PI3K-AKT signaling pathways.^63^ Alternatively, IGF2 may be signaling through insulin receptors (IRs). In the placental vasculature and arterial endothelial cells, insulin acts through IR to activate endothelial nitric oxide synthase (eNOS), which has been proposed to facilitate angiogenesis via HIF-1α, VEGF, and VEGFR-2 ^64^ Indeed, in mice, IGF2 increases labyrinth zone volume and surface area dedicated to transport, ^65^ and perhaps may be why we see that PHF fetal weights are comparable to those of control pregnancies.

We show clear sex-specific changes due to paternal obesity. Indeed, female sex is more likely to be associated with very early preterm preeclampsia,^66^ and in many experimental studies females show compromised placental development after early life adversity,^67, 68^ In our study, females consistently show increased placental transcript levels of angiogenic markers. As we see similar levels of placental blood vessel maturity at term gestation in both males and female placentae, it is possible that females (but not males) require sustained high levels of pro-angiogenic regulators to maintain placental growth and development. This would be consistent with other models of hypoxia where female (but not male) placental labyrinth microstructure was impaired after maternal hypoxia,^69^ suggesting that males somehow compensate in cases of prolonged hypoxia.

### Despite changes in placental vasculature, placental lactogen and maternal glucose metabolism is not altered by paternal obesity

Placental endocrine function, including placental lactogen production by trophoblast giant cells, mediates maternal metabolic adaptation to pregnancy^38^ Two placental lactogens exist in mice encoded by genes *Csh1* (or *Prl3d1*) and *Csh2* (or *Prl3d1*),^28^ PL precipitates maternal insulin resistance facilitating transplacental maternal glucose transfer to the fetus. We found that despite the fact that paternal obesity induced impairments to placental development, maternal glucose tolerance was similar between dams sired by CON and PHF males. Consistent with this observation both the transcript levels of *Csh1* and *Csh2*, and placental giant cell number were similar between groups. These data suggest that paternal obesity-induced changes in placental development may be restricted to vascular impairments amounting to hypoxic environment rather than having effects on maternal pregnancy adaptations.

### Paternal obesity results fetal hepatic ER stress and promotes postnatal metabolic dysfunction

Hepatic ER stress is associated with the developmental of obesity and metabolic complications including the Metabolic Syndrome.^70^ Indeed, *in utero* exposure to nutritional adversity in the form of maternal fructose induces ER stress in fetal liver,^71^ and maternal obesity results in ER stress induction in the fetal gut^4^ and offspring liver.^72^ Although paternal obesity leads to impaired glucose tolerance and liver steatosis in offspring,^73^ no studies have investigated whether changes in the fetal liver, contribute to offspring metabolic dysfunction. We observed increased hepatic transcript levels of key regulators and downstream effectors of the UPR in fetuses of PHF fathers. Paternal obesity increased protein and/or transcript levels of GRP78, PERK and members of the ATF6 (*Edem1*) pathways in fetal livers as well as CHOP and BAX. These increases were accompanied by elevated levels of cleaved caspase-3, suggesting paternal obesity may increase fetal hepatic apoptosis. Although we did not evaluate whether liver steatosis and/or disease, previous work suggests that the UPR is activated in several liver diseases including obesity-associated fatty liver disease.^73^ Thus, prenatal-induced ER stress could precipitate impairments in postnatal hepatic lipid handling, resulting in increased postnatal risk of steatosis.

As ER-stress related liver disease is often associated with changes in hepatic glucose metabolism we investigated transcriptional regulators gluconeogenesis. We found that paternal obesity increased transcript levels of *Ppargclα* and its co-regulator *Hnf4a* (albeit moderate increases in opposing sexes) in fetal livers, As fetal gluconeogenesis is limited due to transplacental transfer of maternal glucose,^74^ these subtle differences may be enhanced postnatally, but this is unknown. Alternatively, increased transcripts of *Hnf4a* in male fetuses could be facilitating hepatocyte differentiation. Recent data suggests that *Hnf4a* may act in concert with a CREB/ATF transcription factor *Crebh* during hepatogenesis and is indirectly stimulated by ER stress.^75^ It is possible that PHF male fetuses compensate, via *Hnf4a-Crebh,* to increase hepatogenesis during *in utero* ER stress, but our study did not test this hypothesis.

We proposed that the functional impacts of these changes in the fetal liver are evident postnatally. We demonstrate that not only is paternal obesity associated with offspring impaired glucose tolerance, but also in whole body energy utilization and metabolism. Previous work has suggested that *in utero* adversity activates the hepatic UPR, increases expression of gluconeogenic genes in pups, and results in glucose intolerance in male offspring.^76^ To our knowledge this is the first time that paternal obesity has been associated with alterations in whole body energetics, including physical activity, heat production, and O_2_ and CO_2_ dynamics. We show that offspring of obese fathers were not only glucose intolerant, but that they ate more and moved less through the day, which may contribute to or be a function of a reduction in their basal metabolic rate. Therefore, this could fuel long-term development of obesity and result in insulin resistance.

We recognize that our study is not without limitations. We did not perform stereology on our placental samples to determine whether the vascular changes we observe in PHF placentae also impact maternal-fetal surface exchange. Additionally, we do not know whether our observations of placental hypoxia result in fetal hypoxia, which could be the driver of hepatic ER stress. Future studies will investigate upstream regulators of placentation and whether alterations in the sperm epigenome of these sires contributed to aberrant placentation.

In conclusion, paternal obesity-induced offspring glucose intolerance and metabolic dysfunction may be driven by impaired placental vascular development, and placental hypoxia. We postulate that impaired placental function results in an adverse intrauterine environment that promotes ER stress-mediated adaptations in fetal hepatic development that predisposes to postnatal impaired energetics and metabolic compromise in offspring.

## ACKNOWLEDGEMENTS

We thank the CNPq Science without Boarders Exchange Program that funded Dr. Ribeiro’s exchange with the Sloboda Lab during the time of data collection. We would like to thank Dr. Gregory R. Steinberg (Department of Medicine, McMaster University) for use of the body composition analyzer and CLAMS and Drs. Rebecca J. Ford and James S. Lally (Department of Medicine, McMaster University) for their assistance with CLAMS analysis. We thank Mr. Andrew De Jong for his assistance with immunoblotting experiments in fetal liver.

P.A.J. is supported by a Thomas Neilson Scholarship, Fred & Helen Knight Enrichment Award, and an Ontario Graduate Scholarship. T.A.R. was supported by CNPq – Conselho Nacional de Desenvolvimento Científico e Tecnológico – Brasil. E.Y. and K.M.K. are supported by the Farncombe Digestive Health Research Institute Student Fellowship. D.M.S. is supported by a Tier 2 Canada Research Chair in Perinatal Programming.

## AUTHOR CONTRIBUTIONS

P.A.J. wrote the manuscript. P.A.J., V.S.P., and T.A.R conducted the animal work, data collection, and data analysis. E.Y. conducted molecular analyses in E18.5 fetal livers. K.M.K. analyzed offspring CLAMS data. J.J.P. conducted IHC experiments in E14.5 and E18.5 placentae. P.A.J., V.S.P., T.A.R., and D.M.S. interpreted the data. D.M.S. designed the experiments and wrote the manuscript. All authors have reviewed and approved the final manuscript.

## REFERENCES

1. Organization WH. Obesity and overweight.). World Health Organization (2020).

2. Fleming TP, et al. Origins of lifetime health around the time of conception: causes and consequences. The Lancet 391, 1842–1852 (2018).

3. Lane M, Robker RL, Robertson SA. Parenting from before conception. Science 345, 756–760 (2014).

4. Gohir W, et al. High-fat diet intake modulates maternal intestinal adaptations to pregnancy and results in placental hypoxia, as well as altered fetal gut barrier proteins and immune markers. J Physiol 597, 3029–3051 (2019).

5. Wallace JG, et al. Obesity During Pregnancy Results in Maternal Intestinal Inflammation, Placental Hypoxia, and Alters Fetal Glucose Metabolism at Mid-Gestation. Sci Rep 9, (2019).

6. Danielzik S, Langnase K, Mast M, Spethmann C, Muller MJ. Impact of parental BMI on the manifestation of overweight 5-7 year old children. Eur J Nutr 41, 132–138 (2002).

7. Ng SF, Lin RC, Laybutt DR, Barres R, Owens JA, Morris MJ. Chronic high-fat diet in fathers programs beta-cell dysfunction in female rat offspring. Nature 467, 963–966 (2010).

8. Fullston T, et al. Paternal obesity initiates metabolic disturbances in two generations of mice with incomplete penetrance to the F2 generation and alters the transcriptional profile of testis and sperm microRNA content. FASEB J 27, 4226–4243 (2013).

9. Binder NK, Hannan NJ, Gardner DK. Paternal diet-induced obesity retards early mouse embryo development, mitochondrial activity and pregnancy health. PLoS One 7, e52304 (2012).

10. Binder NK, Mitchell M, Gardner DK. Parental diet-induced obesity leads to retarded early mouse embryo development and altered carbohydrate utilisation by the blastocyst. Reproduction, Fertility and Development 24, 804–812 (2012).

11. Bakos HW, Mitchell M, Setchell BP, Lane M. The effect of paternal diet-induced obesity on sperm function and fertilization in a mouse model. Int J Androl 34, 402–410 (2011).

12. Chavarro JE, Toth TL, Wright DL, Meeker JD, Hauser R. Body mass index in relation to semen quality, sperm DNA integrity, and serum reproductive hormone levels among men attending an infertility clinic. Fertil Steril 93, 2222–2231 (2010).

13. Hammoud AO, Wilde N, Gibson M, Parks A, Carrell DT, Meikle AW. Male obesity and alteration in sperm parameters. Fertil Steril 90, 2222–2225 (2008).

14. Donkin I, et al. Obesity and Bariatric Surgery Drive Epigenetic Variation of Spermatozoa in Humans. Cell Metab 23, 369–378 (2016).

15. Chen Q, et al. Sperm tsRNAs contribute to intergenerational inheritance of an acquired metabolic disorder. Science 351, 397–400 (2016).

16. Fowden AL, Coan PM, Angiolini E, Burton GJ, Constancia M. Imprinted genes and the epigenetic regulation of placental phenotype. Prog Biophys Mol Biol 106, 281–288 (2011).

17. McPherson NO, et al. When two obese parents are worse than one! Impacts on embryo and fetal development. Am J Physiol Endocrinol Metab 309, E568–581 (2015).

18. Binder NK, Beard SA, Kaitu’u-Lino TJ, Tong S, Hannan NJ, Gardner DK. Paternal obesity in a rodent model affects placental gene expression in a sex-specific manner. Reproduction 149, 435–444 (2015).

19. Mitchell M, et al. Gene expression and epigenetic aberrations in F1-placentas fathered by obese males. Mol Reprod Dev 84, 316–328 (2017).

20. Burton GJ, Fowden AL. The placenta: a multifaceted, transient organ. Philos Trans R Soc Lond B Biol Sci 370, 20140066 (2015).

21. Bar J, Kovo M, Schraiber L, Shargorodsky M. Placental maternal and fetal vascular circulation in healthy non-obese and metabolically healthy obese pregnant women. Atherosclerosis 260, 63–66 (2017).

22. Jauniaux E, Watson AL, Hempstock J, Bao Y, Skepper JN, Burton GJ. Onset of Maternal Arterial Blood Flow and Placental Oxidative Stress. American Journal of Pathology 157, 2111–2122 (2000).

23. Malhotra JD, Kaufman RJ. The endoplasmic reticulum and the unfolded protein response. Semin Cell Dev Biol 18, 716–731 (2007).

24. Westermeier F, Saez PJ, Villalobos-Labra R, Sobrevia L, Farias-Jofre M. Programming of fetal insulin resistance in pregnancies with maternal obesity by ER stress and inflammation. Biomed Res Int 2014, 917672 (2014).

25. Cavallari JF, et al. Muramyl Dipeptide-Based Postbiotics Mitigate Obesity-Induced Insulin Resistance via IRF4. Cell Metab 25, 1063–1074 e1063 (2017).

26. Marcinko K, Sikkema SR, Samaan MC, Kemp BE, Fullerton MD, Steinberg GR. High intensity interval training improves liver and adipose tissue insulin sensitivity. Mol Metab 4, 903–915 (2015).

27. Matthews DR, Hosker JP, Rudenski AS, Naylor BA, Treacher DF, Turner RC. Homeostasis model assessment: insulin resistance and beta-cell function from fasting plasma glucose and insulin concentrations in man. Diabetologia 28, 412–419 (1985).

28. Yamaguchi M, Ogren L, Endo H, Thordarson G, Bigsby RM, Talamantes F. Production of mouse placental lactogen-I and placental lactogen-II by the same giant cell. Endocrinology 131, 1595–1602 (1992).

29. Krock BL, Skuli N, Simon MC. Hypoxia-induced angiogenesis: good and evil. Genes Cancer 2, 1117–1133 (2011).

30. Hu X, et al. Phosphoinositide 3-Kinase (PI3K) Subunit p110δ Is Essential for Trophoblast Cell Differentiation and Placental Development in Mouse. Scientific Reports 6, (2016).

31. Iwawaki T, Akai R, Yamanaka S, Kohno K. Function of IRE1 alpha in the placenta is essential for placental development and embryonic viability. PNAS 106, 16657–16662 (2009).

32. Charnock-Jones DS. Placental hypoxia, endoplasmic reticulum stress and maternal endothelial sensitisation by sFLT1 in pre-eclampsia. J Reprod Immunol 114, 81–85 (2016).

33. Smith JA. Regulation of Cytokine Production by the Unfolded Protein Response; Implications for Infection and Autoimmunity. Frontiers in Immunology 9, (2018).

34. Sferruzzi-Perri AN, Sandovici I, Constancia M, Fowden AL. Placental phenotype and the insulin-like growth factors: resource allocation to fetal growth. J Physiol 595, 5057–5093 (2017).

35. Soubry A, et al. Paternal obesity is associated with IGF2hypomethylation in newborns-results from a Newborn Epigenetics Study (NEST) cohort. BMC Medicine 11, (2013).

36. Wu J, Zhu AX. Targeting insulin-like growth factor axis in hepatocellular carcinoma. Journal of Hematology & Oncology 4, (2011).

37. Street ME, Viani I, Ziveri MA, Volta C, Smerieri A, Bernasconi S. Impairment of insulin receptor signal transduction in placentas of intra-uterine growth-restricted newborns and its relationship with fetal growth. European Journal of Endocrinology 164, 45–52 (2011).

38. Newbern D, Freemark M. Placental hormones and the control of maternal metabolism and fetal growth. Curr Opin Endocrinol Diabetes Obes 18, 409–416 (2011).

39. Campbell JM, Lane M, Owens JA, Bakos HW. Paternal obesity negatively affects male fertility and assisted reproduction outcomes: a systematic review and meta-analysis. Reprod Biomed Online 31, 593–604 (2015).

40. Pearce KL, Hill A, Tremellen KP. Obesity related metabolic endotoxemia is associated with oxidative stress and impaired sperm DNA integrity. Basic Clin Androl 29, 6 (2019).

41. Lau HH, Ng NHJ, Loo LSW, Jasmen JB, Teo AKK. The molecular functions of hepatocyte nuclear factors-In and beyond the liver. J Hepatol 68, 1033–1048 (2018).

42. Park S, Jang A, Bouret SG. Maternal obesity-induced endoplasmic reticulum stress causes metabolic alterations and abnormal hypothalamic development in the offspring. PLoS Biol 18, e3000296 (2020).

43. Perez-Arancibia R, Rivas A, Hetz C. (off)Targeting UPR signaling: the race toward intervening ER proteostasis. Expert Opin Ther Targets 22, 97–100 (2018).

44. Soares Moretti AI, Martins Laurindo FR. Protein disulfide isomerases: Redox connections in and out of the endoplasmic reticulum. Archives of Biochemistry and Biophysics 617, 106–119 (2017).

45. Speakman JR. Measuring energy metabolism in the mouse-theoretical, practical, and analytical considerations. Front Physiol 4, 34 (2013).

46. Aljunaidy MM, Morton JS, Cooke CM, Davidge ST. Prenatal hypoxia and placental oxidative stress: linkages to developmental origins of cardiovascular disease. Am J Physiol Regul Integr Comp Physiol 313, R395–R399 (2017).

47. Ahn H, Park J, Gilman-Sachs A, Kwak-Kim J. Immunologic characteristics of preeclampsia, a comprehensive review. Am J Reprod Immunol 65, 377–394 (2011).

48. Zhao H, Narasimhan P, Kalish F, Wong RJ, Stevenson DK. Dysregulation of hypoxia-inducible factor-1 (Hif1) expression in the Hmox1-deficient placenta. Placenta 99, 108–116 (2020). ^α^

49. Ghosh R, et al. Transcriptional regulation of VEGF-A by the unfolded protein response pathway. PLoS One 5, e9575 (2010).

50. Pereira ER, Frudd K, Awad W, Hendershot LM. Endoplasmic reticulum (ER) stress and hypoxia response pathways interact to potentiate hypoxia-inducible factor 1 (HIF-1) transcriptional activity on targets like vascular endothelial growth factor (VEGF). J Biol Chem 289, 3352–3364 (2014).

51. Choi SH, Park JY. Erratum to: Regulation of the hypoxic tumor environment in hepatocellular carcinoma using RNA interference. Cancer Cell Int 17, 69 (2017).

52. Tam AB, Mercado EL, Hoffmann A, Niwa M. ER stress activates NF-kappaB by integrating functions of basal IKK activity, IRE1 and PERK. PLoS One 7, e45078 (2012).

53. Eltzschig HK, Carmeliet P. Hypoxia and inflammation. N Engl J Med 364, 656–665 (2011).

54. Baun M, Hay-Schmidt A, Edvinsson L, Olesen J, Jansen-Olesen I. Pharmacological characterization and expression of VIP and PACAP receptors in isolated cranial arteries of the rat. Eur J Pharmacol 670, 186–194 (2011).

55. Moore DA, et al. Hypoxia-inducible factor-1 Regulates Matrix Metalloproteinase-14 Expression: Underlying Effects of Hypoxia and Statins. Heart 100, A111–A112 (2014).

56. Chen J, Khalil RA. Matrix Metalloproteinases in Normal Pregnancy and Preeclampsia. Prog Mol Biol Transl Sci 148, 87–165 (2017).

57. Zhang Z, Grewer C. The sodium-coupled neutral amino acid transporter SNAT2 mediates an anion leak conductance that is differentially inhibited by transported substrates. Biophys J 92, 2621–2632 (2007).

58. Manta-Vogli PD, Schulpis KH, Dotsikas Y, Loukas YL. The significant role of amino acids during pregnancy: nutritional support. J Matern Fetal Neonatal Med 33, 334–340 (2020).

59. Menchini RJ, Chaudhry FA. Multifaceted regulation of the system A transporter Slc38a2 suggests nanoscale regulation of amino acid metabolism and cellular signaling. Neuropharmacology 161, 107789 (2019).

60. Brocker C, Thompson DC, Vasiliou V. The role of hyperosmotic stress in inflammation and disease. BioMolecular Concepts 3, 345–364 (2012).

61. Nishimura T, et al. System A amino acid transporter SNAT2 shows subtype-specific affinity for betaine and hyperosmotic inducibility in placental trophoblasts. Biochimica et Biophysica Acta (BBA) - Biomembranes 1838, 1306–1312 (2014).

62. Morotti M, et al. Hypoxia-induced switch in SNAT2/SLC38A2 regulation generates endocrine resistance in breast cancer. Proc Natl Acad Sci U S A 116, 12452–12461 (2019).

63. Forbes K, Westwood M, Baker PN, Aplin JD. Insulin-like growth factor I and II regulate the life cycle of trophoblast in the developing human placenta. American Journal of Physiology-Cell Physiology 294, C1313–C1322 (2008).

64. Lassance L, et al. Hyperinsulinemia Stimulates Angiogenesis of Human Fetoplacental Endothelial Cells: A Possible Role of Insulin in Placental Hypervascularization in Diabetes Mellitus. The Journal of Clinical Endocrinology & Metabolism 98, E1438–E1447 (2013).

65. Sferruzzi-Perri AN, Owens JA, Pringle KG, Robinson JS, Roberts CT. Maternal insulin-like growth factors-I and -II act via different pathways to promote fetal growth. Endocrinology 147, 3344–3355 (2006).

66. Taylor BD, et al. The impact of female fetal sex on preeclampsia and the maternal immune milieu. Pregnancy Hypertens 12, 53–57 (2018).

67. Osei-Kumah A, Smith R, Jurisica I, Caniggia I, Clifton VL. Sex-specific differences in placental global gene expression in pregnancies complicated by asthma. Placenta 32, 570–578 (2011).

68. O’Connell BA, Moritz KM, Roberts CT, Walker DW, Dickinson H. The placental response to excess maternal glucocorticoid exposure differs between the male and female conceptus in spiny mice. Biol Reprod 85, 1040–1047 (2011).

69. Cuffe JSM, et al. Mid-to late term hypoxia in the mouse alters placental morphology, glucocorticoid regulatory pathways and nutrient transporters in a sex-specific manner. The Journal of physiology 592, 3127–3141 (2014).

70. Hotamisligil GS. Endoplasmic reticulum stress and the inflammatory basis of metabolic disease. Cell 140, 900–917 (2010).

71. Rodrigo S, et al. Effects of Maternal Fructose Intake on Perinatal ER-Stress: A Defective XBP1s Nuclear Translocation Affects the ER-stress Resolution. Nutrients 11, 1935 (2019).

72. de Almeida-Faria J, et al. Maternal obesity during pregnancy leads to adipose tissue ER stress in mice via miR-126-mediated reduction in Lunapark. Diabetologia 64, 890–902 (2021).

73. Ornellas F, Souza-Mello V, Mandarim-de-Lacerda CA, Aguila MB. Programming of obesity and comorbidities in the progeny: lessons from a model of diet-induced obese parents. PLoS One 10, e0124737 (2015).

74. Kalhan S, Parimi P. Gluconeogenesis in the fetus and neonate. Semin Perinatol 24, 94–106 (2000).

75. Luebke-Wheeler J, et al. Hepatocyte nuclear factor 4alpha is implicated in endoplasmic reticulum stress-induced acute phase response by regulating expression of cyclic adenosine monophosphate responsive element binding protein H. Hepatology 48, 1242–1250 (2008).

76. Deodati A, et al. The exposure to uteroplacental insufficiency is associated with activation of unfolded protein response in postnatal life. PLoS One 13, e0198490 (2018).

