## Supplemental Figure S1 for "Paternal obesity results in placental hypoxia and sex-specific impairments in placental vascularization and offspring metabolic function"

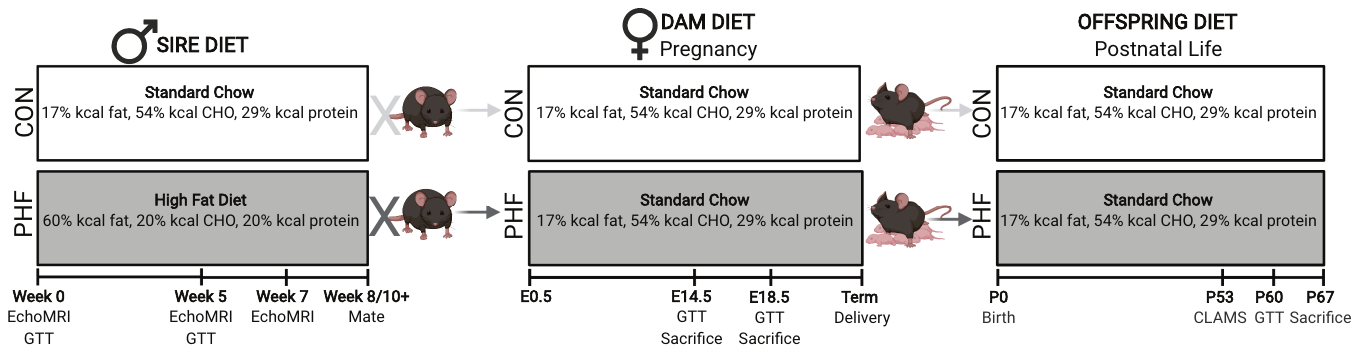

**Supplemental Figure S1. Experimental Design.**

C57BL/6J male mice were assigned to receive either a standard chow diet (control, CON, Harlan 8640 Teklad 2/5 Rodent Diet; n = 49) or high fat diet (paternal high fat, PHF; Diets Inc. D12492; n = 61) *ad libitum* for 8-10 weeks. Males were subjected to an echoMRI at weeks 0, 5, and 7 of diet and a glucose tolerance test (GTT) at weeks 0, and 5 of diet. After 8-10 weeks on their respective diets, CON and PHF males were time-mated with C57BL/6J female mice to generate pregnancies and offspring. Female mice were fed a standard chow diet (Harlan 8640 Teklad 22/5 Rodent Diet). A subset of pregnant females was subjected to a GTT to assess glucose tolerance at mid-gestation (embryonic day (E) 14.5) or term gestation (E18.5). Another subset of pregnant females was sacrificed at E14.5 or E18.5. Placentae and fetal liver were collected for histological and molecular analyses. A third subset of pregnant females was allowed to deliver pups. Male and female CON and PHF offspring were subjected to Comprehensive Lab Animal Monitoring System (CLAMS) for metabolic profiling at postnatal day P53, then to a standard GTT to assess glucose tolerance at P60, and subsequently sacrificed at P67.
