## Supplemental Figure S2 for "Paternal obesity results in placental hypoxia and sex-specific impairments in placental vascularization and offspring metabolic function"

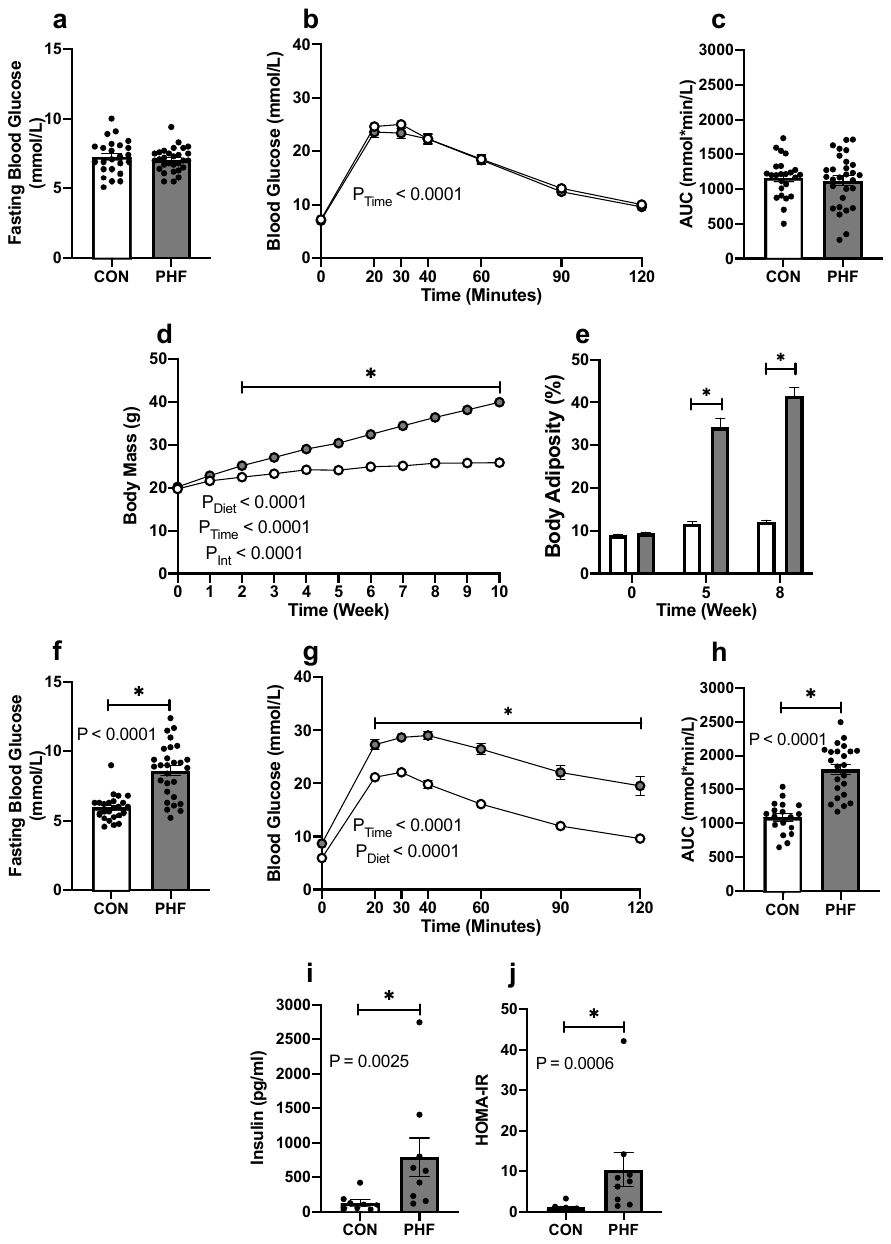

**Supplemental Figure S2. Male mice consuming a high fat diet become obese, with increased body mass and body adiposity, hyperglycemia, and glucose intolerance.**

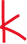

Male mice were assigned to either: 1) Control (CON; n = 49): fed a standard chow diet *ad libitum*; or 2) Paternal High Fat (PHF; n = 61): fed a high fat (HF) diet *ad libitum* for ten weeks. Males were subjected to an echoMRI and glucose tolerance test at 0 (baseline), 5, and 7 weeks after diet consumption. (**a**) Prior to diet allocation at baseline, CON and PHF males had similar fasting blood glucose levels and (**b**) glucose tolerance. (**c**) PHF males were significantly heavier after consuming a HF diet for two weeks. (**d**) PHF body adiposity was increased at 5 and (**e**) 7 weeks of diet. (**f**) After 5 weeks of diet, PHF males had higher fasting blood glucose levels, (**g**) were glucose intolerant, (**h**) with an elevated area under the curve (AUC), (**i**) had higher serum insulin concentrations, and (**j**) HOMA-IR. Data are presented as mean ± SEM. Student’s t-test or Mann-Whitney U test, two-way ANOVA, or two-way repeated measures ANOVA with main effects of paternal diet and time as factors. **P* < 0.05 using Bonferroni’s post-hoc for multiple comparison, where appropriate. CON = Control (open circles and bars); PHF = Paternal high fat diet-induced obesity (grey circles and bars).
