## Supplemental Figure S3 for "Paternal obesity results in placental hypoxia and sex-specific impairments in placental vascularization and offspring metabolic function"

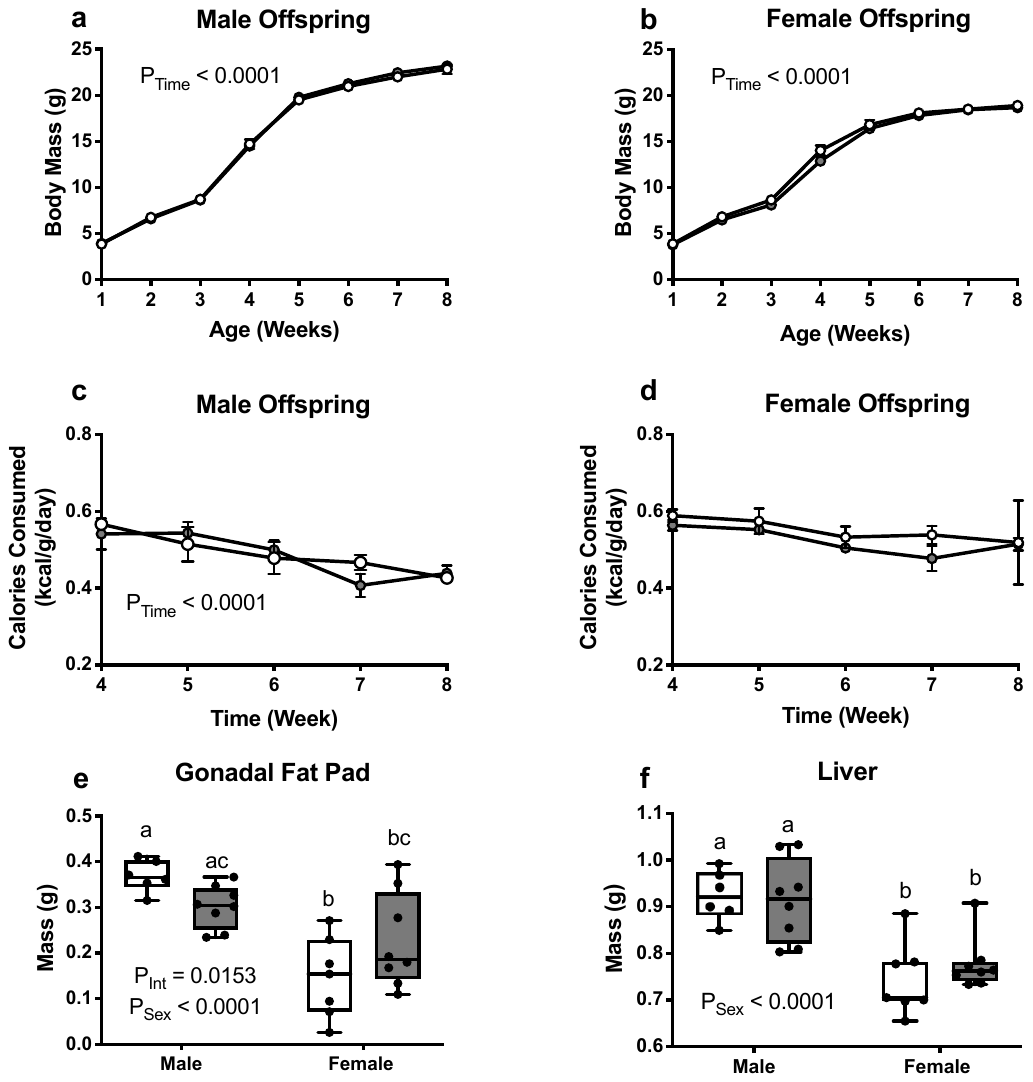

**Supplemental Figure S3. Offspring of obese fathers do not show an appreciable postnatal anthropometric phenotype.**

Female mice were time-mated with CON or PHF male mice to generate offspring that were sacrificed as young adults at postnatal day 67 (P67). (**a**) Body mass in male and (**b**) female offspring. (**c**) Postnatal daily caloric intake of male and (**d**) female offspring. (**e**) Gonadal fat pad mass, and (**f**) liver mass at P67 in CON and PHF offspring. CON; n = 6, PHF; n = 8. Data are presented as mean ± SEM or as box plots; min to max with line representing the median. Two-way ANOVA with main effects of paternal diet and offspring sex as factors or two-way repeated measures ANOVA with main effects of paternal diet and time as factors using Bonferroni’s post-hoc for multiple comparison. Box plots with different letters indicate significance *P* < 0.05. CON = Control (open circles and box plots); PHF = Paternal high fat diet-induced obesity (grey circles and box plots).
