## Supplemental Table S1 for "Paternal obesity results in placental hypoxia and sex-specific impairments in placental vascularization and offspring metabolic function"

**Supplemental Table S1.** Primer sequences.

| **Gene^a^** | **Forward Primer Sequence** | **Reverse Primer Sequence** | **GenBank Accession Number** |
| --- | --- | --- | --- |
| *Atf4* | GGGTTCTGTCTTCCACTCCA | AAGCAGCAGAGTCAGGCTTTC | NM_009716.3 |
| *Atf6* | AGAAAGCCCGCATTCTCCAG | CTCACTCCCAGAATTCCTACTGAT | NM_001081304.1 |
| *β-actin* | AGATCAAGATCATTGCTCCT | ACGCAGCTCAGTAACAGTC | NM_007393.5 |
| *Bax* | GATCCAAGACCAGGGTGGCT* | CCTTCCCCCATTCATCCCAG* | NM_007527.3 |
|  | TGCAGAGGATGATTGCTGACG^#^ | CCAGCCACCCTGGTCTTG^#^ |  |
| *Bcl2* | CACTCTGGGTGCATACCTGG | GTTTGGGGCAGGTTTGTCG | NM_009741.5 |
| *B2m* | CTCGGTGACCCTGGTCTTTC | TTGAGGGGTTTTCTGGATAGCA | NM_009735.3 |
| *Csh1* | TGGAGCCTACATTGTGGTGGA | CATTCCTGCGGAGCCTGAAAG | NM_001205322.1 |
| *Csh2* | GTCCACCAGACAACATCGGA | CTGCTGCCACCATGTGTTTC | NM_008865.3 |
| *Ddit3* | CCACCACACCTGAAAGCAGAA | AGGTGAAAGGCAGGGACTCA | NM_007837.4 |
| *Edem1* | CTACCTGCGAAGAGGCCG | GTTCATGAGCTGCCCACTGA | NM_138677.2 |
| *Fabp4* | AAGCTGGTGGTGGAATGTGTTA | CCTCTTCCTTTGGCTCATGC | NM_024406.3 |
| *Hmox1* | AGGCTTTAAGCTGGTGATGGC | GGGGCATAGACTGGGTTCTG | NM_010442.2 |
| *Hnf4a* | GGGTAGGGGAGAATGCGACT | TTCAGATGGGGACGTGTCATT | NM_008261.3 |
| *Hprt* | CAGTCCCAGCGTCGTGATTA | TCGAGCAAGTCTTTCAGTCCT | NM_013556.2 |
| *Hspa5* | TTCAGCCAATTATCAGCAAACTCT | TTTTCTGATGTATCCTCTTCACCAGT | NM_001163434.1 |
| *G6pc* | CTGTCCCGGATCTACCTTGC | CACAGCAATGCCTGACAAGAC | NM_008061.4 |
| *Igf2* | CCAGCCCTAAGATACCCTAAAGA | GAAGCACCAACATCGACTTCC | NM_010514.3 |
| *Il1b* | GCCACCTTTTGACAGTGATGAG | GACAGCCCAGGTCAAAGGTT | NM_008361.4 |
| *Il6* | GGGACTGATGCTGGTGACAA | ACAGGTCTGTTGGGAGTGGT | NM_001314054.1 |
| *Ipo8* | AGACGGAGCTTAACCAGTCCT | GCAAGCTGGGGGCAAAAT | NM_001081113.1 |
| *Irs1* | GGACATCACAGCAGAATGAAGAC | AGACGTGAGGTCCTGGTTGT | NM_010570.4 |
| *Irs2* | TCCAGGCACTGGAGCTTTG | CTTCACTCTTTCACGACTGTGG | NM_001081212.2 |
| *Mmp2* | AACGGTCGGGAATACAGCAG | GGTAAACAAGGCTTCATGGGG | NM_008610.3 |
| *Mmp14* | GATAAGCCCAAAAACCCCGC | AACCATCGCTCCTTGAAGACA | NM_008608.4 |
| *Nono* | GCCAGAATGAAGGCTTGACTAT | TATCAGGGGGAAGATTGCCCA | NM_001252518.1 |
| *Pck1* | GGGTGGAAGGTCGAATGTGT | TAGCCCTTAAGTTGCCTTGGG | NM_011044.3 |
| *Pdia2* | GTCCCGCTTCCTCGTCATAC | CCATTACGAACAACACCTGCC | NM_001081070.1 |
| *Ppargc1a* | TTGACTGGCGTCATTCGGG | AGAGCAGCACACTCTATGTCAC | NM_008904.2 |
| *Ppia* | CTTCGAGCTGTTTGCAGACA | TGGCGTGTAAAGTCACCAC | NM_008907.1 |
| *Ppp1r15a* | CCAGCGTTGTCTACCAGGAG | AGTGTACCTTCCGAGCTTTTAGA | NM_008654.2 |
| *Slc2a1* | GGCTTGCTTGTAGAGTGACG | TGTAGAACTCCTCAATAACCTTCTG | NM_011400.3 |
| *Slc2a3* | AATAGGTAGGCTGGGCTTCG | AGAGATGGGGTCACCTTCGTT | NM_011401.4 |
| *Slc38a2* | GCAGTGGAATCCTTGGGCTT | TAAAGATCCTCCTTCGTTGGCAG | NM_175121.3 |
| *Tnf* | TAGCCACGTCGTAGCAAAC | ACAAGGTACAACCCATCGGC | NM_013693.3 |
| *Traf6* | GCACGGAAACTTGGGTCTT | CTCTGTTGTCAGTCGACTTG | NM_009424.3 |
| *Xbp1s* | CTGAGTCCGAATCAGGTGCAG | GTCCATGGGAAGATGTTCTGG | NM_001271730.1 |
| *Xbp1t* | TGGCCGGGTCTGCTGAGTCCG | GTCCATGGGAAGATGTTCTGG | NM_001271730.1 |
| *Ywhag* | GTGACCGAGCTGAACGAAC | GATGCTCCTGATGACCCTCC | NM_018871.3 |
| *Ywhaz* | TGTGTCTCCCAATGAAAGCTCTA | GACGTCAAACGCTTCTGGCT | NM_011740.3 |

^a^*Atf4,* activating transcription factor 4; *Atf6,* activating transcription factor 6; *Bax,* bcl2 associated x, apoptosis regulator; *Bcl2,* bcl2 apoptosis regulator; *β-actin,* beta-actin; *B2m,* beta-2-microglobulin; *Csh1*, chorionic somatomammotropin hormone 1; *Csh2*, chorionic somatomammotropin hormone 2; *Ddit3,* DNA damage inducible transcript 3; *Edem1,* ER degradation enhancing alpha-mannosidase like protein 1; *Fabp4,* fatty acid binding protein 4; *G6pc*, glucose-6-phosphatase, catalytic; *Hmox1,* heme oxygenase 1; *Hnf4a*, hepatocyte nuclear factor 4 alpha; *Hprt*, hypoxanthine guanine phosphoribosyltransferase; *Hspa5,* heat shock protein family a member 5; *Igf2*, insulin like growth factor 2; *Il1b*, interleukin 1 beta; *Il6*, interleukin 6; *Ipo8,* importin 8; *Irs1*, insulin receptor substrate 1; *Irs2*, insulin receptor substrate 2; *Mmp2,* matrix metallopeptidase 2; *Mmp14,* matrix metallopeptidase 14; *Nono,* non-POU domain containing octamer binding; *Pck1*, phosphoenolpyruvate carboxykinase 1, cytosolic; *Pdia2*, protein disulfide isomerase family a member 2; *Ppargc1a*, peroxisome proliferative activated receptor gamma, coactivator 1 alpha; *Ppia,* peptidylprolyl isomerase a; *Ppp1r15a,* protein phosphatase 1 regulatory subunit 15a; *Slc2a1,* solute carrier family 2 member 1; *Slc2a3,* solute carrier family 2 member 3; *Slc38a2,* solute carrier family 38 member 2; *Tnf,* tumor necrosis factor; *Traf6,* TNF receptor associated factor 6; *Xbp1s*, X-box binding protein 1 (spliced); *Xbp1t*, X-box binding protein 1 (total); *Ywhag,* tyrosine 3-monooxygenase/tryptophan 5-monooxygenase activation protein gamma; Ywhaz, tyrosine 3-monooxygenase/tryptophan 5-monooxygenase activation protein zeta.

*Primer sequence used for placental qPCR assays.

^#^Primer sequence used for E18.5 fetal liver qPCR assays.
