## Supplemental Table S2 for "Paternal obesity results in placental hypoxia and sex-specific impairments in placental vascularization and offspring metabolic function"

**Supplemental Table S2.** Housekeeping genes used to normalize quantitative PCR data.

| **Tissue** | **Time Point** | **Housekeeping Genes^a^** |
| --- | --- | --- |
| Placenta | E14.5 | *β-actin*, *B2m*, *Nono*, *Ppia*, and *Ywhag* |
|  | E18.5 | *Hprt, and Ipo8* |
| Liver | E14.5 | *B2m*, *Nono*, and *Ywhag* |
|  | E18.5 | *Ipo8*, and *Ywhaz* |

^a^*β-actin*, beta-actin; *B2m,* beta-2-microglobulin; *Hprt*, hypoxanthine guanine phosphoribosyltransferase; *Ipo8,* importin 8; *Nono,* non-POU domain containing octamer binding; *Ppia,* peptidylprolyl isomerase a; *Ywhag,* tyrosine 3-monooxygenase/tryptophan 5-monooxygenase activation protein gamma; Ywhaz, tyrosine 3-monooxygenase/tryptophan 5-monooxygenase activation protein zeta.
