## Supplemental Table S3 for "Paternal obesity results in placental hypoxia and sex-specific impairments in placental vascularization and offspring metabolic function"

**Supplemental Table S3.** Antibodies used for placental immunohistochemical staining.^a^

| **Antibody Name** | **Target** | **Product Number** | **Dilution** | **Source** |
| --- | --- | --- | --- | --- |
| α-Actin (1A4) Antibody | α-SMA | sc-32251 | 1:400 | Santa Cruz Biotechnology |
| Anti-Carbonic Anhydrase 9/CA9 Antibody | CA IX | ab15086 | 1:600 | Abcam |
| Anti-CD31 Antibody [P2B1] | CD31 | ab24590 | 1:200 | Abcam |
| VEGF (A-20) Antibody | VEGF-A | sc-152 | 1:400 | Santa Cruz Biotechnology |
| VEGFR2 (A-3) Antibody | VEGFR-2 | sc-6251 | 1:400 | Santa Cruz Biotechnology |

^a^α-SMA, alpha-smooth muscle actin; CA IX, carbonic anhydrase; CD31, cluster of differentiation 31; VEGF-A, vascular endothelial growth factor A; VEGFR-2, vascular endothelial growth factor receptor 2.
