## Supplemental Table S4 for "Paternal obesity results in placental hypoxia and sex-specific impairments in placental vascularization and offspring metabolic function"

**Supplemental Table S4.** Primary antibodies used for immunoblotting.^a^

| **Antibody Name** | **Protein** | **Product Number** | **Dilution** | **Host** | **Source** |
| --- | --- | --- | --- | --- | --- |
| Anti-GRP78 BiP Antibody | GRP78 | ab21685 | 1:2000 | Rabbit | Abcam |
| Phospho-PERK (Thr980) Monoclonal Antibody | Phospho-PERK | MA-15033 | 1:500 | Rabbit | ThermoFisher |
| PERK (C33E10) Rabbit mAb | PERK | 3192 | 1:1000 | Rabbit | Cell Signaling Technology® |
| Phospho-eIF2α (Ser51) Antibody | Phospho-eIF2α | 9721 | 1:1000 | Rabbit | Cell Signaling Technology® |
| Anti-IRE1 Antibody | IRE1α | ab37073 | 1:1000 | Rabbit | Abcam |
| Phospho-IRE1 (Phospho S724) Antibody | Phospho-IRE1α | ab48187 | 1:1000 | Rabbit | Abcam |
| eIF2α Antibody | eIF2α | 9722 | 1:1000 | Rabbit | Cell Signaling Technology® |
| Cleaved Caspase-3 (Asp175) Antibody | Cleaved CASP3 | 9661 | 1:1000 | Rabbit | Cell Signaling Technology® |
| Caspase-3 Antibody | CASP3 | 9662 | 1:1000 | Rabbit | Cell Signaling Technology® |
| HIF-1 alpha Antibody | HIF-1α | NB100-479 | 1:1000 | Rabbit | Novus Biologicals |
| β-actin (13E5) Rabbit mAb (HRP Conjugate)^b^ | β-actin | 5125 | 1:5000 | Rabbit | Cell Signaling Technology® |
| TBP Antibody^bc^ | TBP | 8515 | 1:1000 | Rabbit | Cell Signaling Technology® |

^a^Asp175, aspartic acid 175; BiP, binding immunoglobulin protein; eIF2α, eukaryotic initiation factor 2 alpha; GRP78, glucose-regulated protein 78; HIF-1, hypoxia-inducible factor 1; HRP, horseradish perodixase; IRE1, inositol requiring enzyme 1; mAb, monoclonal antibody; PERK, protein kinase RNA-like endoplasmic reticulum kinase; Ser51, serine 51; S724, serine 724; TBP, TATA-binding protein.

^b^Primary antibody used as positive internal control.

^c^Primary antibody used to immunostain nuclear protein extracts only.
