## Supplemental Table S5 for "Paternal obesity results in placental hypoxia and sex-specific impairments in placental vascularization and offspring metabolic function"

**Supplementary Table S5.** Activity levels and relative transcript levels of downstream IRE1 pro-inflammatory signaling genes in mid-gestation (E14.5) and term gestation (E18.5) placentae.*

| **Activity Levels/*Gene* transcript levels** | **Relative Transcript Levels** | | | | **Main Effect** |
| --- | --- | --- | --- | --- | --- |
|  | **Sire Group** | | | |  |
|  | **CON** | | **PHF** | |  |
|  | **Fetal Sex** | | **Fetal Sex** | |  |
|  | Male  (n = 8-9) | Female  (n = 9) | Male  (n = 5-8) | Female  (n = 6-8) |  |
| **E14.5** | | | | | |
| NF-κB activity | 0.169 ± 0.002 | 0.173 ± 0.003 | 0.173 ± 0.002 | 0.171 ± 0.006 | P_Diet_ = 0.7462 |
|  |  |  |  |  | P_Sex_ = 0.8483 |
|  |  |  |  |  | P_Int_ = 0.4041 |
| *Traf6* | 0.888 ± 0.037 | 0.920 ± 0.079 | 0.830 ± 0.070 | 1.007 ± 0.020 | P_Diet_ = 0.8149 |
|  |  |  |  |  | P_Sex_ = 0.1067 |
|  |  |  |  |  | P_Int_ = 0.2584 |
| *Tnf* | 0.868 ± 0.089 | 1.309 ± 0.197 | 0.950 ± 0.117 | 0.981 ± 0.164 | P_Diet_ = 0.4129 |
|  |  |  |  |  | P_Sex_ = 0.1231 |
|  |  |  |  |  | P_Int_ = 0.1774 |
| *Il1b* | 1.128 ± 0.093^ab^ | 1.470 ± 0.142^a^ | 0.868 ± 0.079^b^ | 1.614 ± 0.192^b^ | P_Diet_ = 0.6708 |
|  |  |  |  |  | **P_Sex_ = 0.0004** |
|  |  |  |  |  | P_Int_ = 0.1493 |
| *Il6* | 0.882 ± 0.082 | 0.925 ± 0.143 | 1.023 ± 0.109 | 0.906 ± 0.085 | P_Diet_ = 0.5857 |
|  |  |  |  |  | P_Sex_ = 0.7438 |
|  |  |  |  |  | P_Int_ = 0.4755 |
| **E18.5** | | | | | |
| NF-κB activity | 0.140 ± 0.016^a^ | 0.207 ± 0.017^a^ | 0.159 ± 0.023^a^ | 0.184 ± 0.036^a^ | P_Diet_ = 0.9366 |
|  |  |  |  |  | **P_Sex_ = 0.0492** |
|  |  |  |  |  | P_Int_ = 0.3440 |
| *Traf6* | 0.595 ± 0.059 | 0.622 ± 0.074 | 0.576 ± 0.039 | 0.725 ± 0.070 | P_Diet_ = 0.5543 |
|  |  |  |  |  | P_Sex_ = 0.2182 |
|  |  |  |  |  | P_Int_ = 0.3869 |
| *Tnf* | 1.394 ± 0.291 | 1.386 ± 0.190 | 1.121 ± 0.112 | 0.850 ± 0.107 | P_Diet_ = 0.0833 |
|  |  |  |  |  | P_Sex_ = 0.5398 |
|  |  |  |  |  | P_Int_ = 0.5623 |
| *Il1b* | 0.692 ± 0.134 | 1.033 ± 0.162 | 0.976 ± 0.217 | 1.037 ± 0.185 | P_Diet_ = 0.4179 |
|  |  |  |  |  | P_Sex_ =0.2602 |
|  |  |  |  |  | P_Int_ = 0.4296 |
| *Il6* | 0.731 ± 0.080^a^ | 0.635 ± 0.088^a^ | 0.939 ± 0.206^a^ | 1.989 ± 0.845^a^ | **P_Diet_ = 0.0431** |
|  |  |  |  |  | P_Sex_ = 0.2049 |
|  |  |  |  |  | P_Int_ = 0.1309 |

CON, placentae from litters generated by CON sires; HF, placentae from litters generated by HF-fed sires; *Il1b*, interleukin 1 beta; *Il6*, interleukin 6; Int, interaction, NF-κB, nuclear factor-kappa B; *Tnf,* tumor necrosis factor; *Traf6,* TNF receptor associated factor 6.

*Data are presented as mean ± SEM. Two-way ANOVA with main effects of sire diet and fetal sex as factors using Bonferroni’s post-hoc for multiple comparisons. Groups that do not share the same letter indicate significance *P* < 0.05.
