## Supplemental Table S6 for "Paternal obesity results in placental hypoxia and sex-specific impairments in placental vascularization and offspring metabolic function"

**Supplemental Table S6.** Protein and relative transcript levels of PERK signaling factors in mid-gestation (E14.5) and term gestation (E18.5) placentae.*

| **Protein/Gene** | **Protein Levels or Relative Transcript Levels** | | | | **Main Effect** |
| --- | --- | --- | --- | --- | --- |
|  | **Sire Group** | | | |  |
|  | **CON** | | **PHF** | |  |
|  | **Fetal Sex** | | **Fetal Sex** | |  |
|  | Male  (n = 8-9) | Female  (n = 8-9) | Male  (n = 6-8) | Female  (n = 6-8) |  |
| **E14.5** | | | | | |
| Phospho-eIF2α: Total eIF2α | 1.024 ± 0.116 | 1.023 ± 0.086 | 1.204 ± 0.097 | 1.104 ± 0.189 | P_Diet_ = 0.3195 |
|  |  |  |  |  | P_Sex_ = 0.7009 |
|  |  |  |  |  | P_Int_ = 0.7032 |
| *Atf4* | 1.001 ± 0.072 | 1.157 ± 0.088 | 1.115 ± 0.112 | 1.347 ± 0.235 | P_Diet_ = 0.2754 |
|  |  |  |  |  | P_Sex_ = 0.1655 |
|  |  |  |  |  | P_Int_ = 0.7824 |
| *Ddit3* | 0.916 ± 0.050 | 0.997 ± 0.087 | 0.861 ± 0.076 | 0.869 ± 0.085 | P_Diet_ = 0.2346 |
|  |  |  |  |  | P_Sex_ = 0.5586 |
|  |  |  |  |  | P_Int_ = 0.6365 |
| **E18.5** | | | | | |
| Phospho-eIF2α: Total eIF2α | 1.100 ± 0.176 | 1.045 ± 0.148 | 1.295 ± 0.432 | 0.892 ± 0.327 | P_Diet_ = 0.9360 |
|  |  |  |  |  | P_Sex_ = 0.3863 |
|  |  |  |  |  | P_Int_ = 0.5085 |
| *Atf4* | 1.019 ± 0.090 | 1.079 ± 0.140 | 1.120 ± 0.107 | 1.188 ± 0.193 | P_Diet_ = 0.4456 |
|  |  |  |  |  | P_Sex_ = 0.6391 |
|  |  |  |  |  | P_Int_ = 0.9779 |
| *Ddit3* | 1.213 ± 0.210 | 1.085 ± 0.175 | 0.940 ± 0.069 | 0.983 ± 0.167 | P_Diet_ = 0.3186 |
|  |  |  |  |  | P_Sex_ = 0.8189 |
|  |  |  |  |  | P_Int_ = 0.6468 |

CON, placentae from litters generated by CON sires; PHF, placentae from litters generated by HF-fed sires; *Atf4,* activating transcription factor 4;

*Ddit3,* DNA damage inducible transcript 3; eIF2α, eukaryotic initiation transcription factor 2A; Int, interaction.

*Data are presented as mean ± SEM. Two-way ANOVA with main effects of sire diet and fetal sex as factors using Bonferroni’s post-hoc for multiple comparisons. Groups that do not share the same letter indicate significance *P* < 0.05.
