## Supplemental Table S7 for "Paternal obesity results in placental hypoxia and sex-specific impairments in placental vascularization and offspring metabolic function"

**Supplemental Table S7.** Relative transcript levels of nutrient transporters in mid-gestation (E14.5) and term gestation (E18.5) placentae.*

| **Gene** | **Protein Levels or Relative Transcript Levels** | | | | **Main Effect** |
| --- | --- | --- | --- | --- | --- |
|  | **Sire Group** | | | |  |
|  | **CON** | | **PHF** | |  |
|  | **Fetal Sex** | | **Fetal Sex** | |  |
|  | Male  (n = 8-9) | Female  (n = 9) | Male  (n = 6-8) | Female  (n = 5-8) |  |
| **E14.5** | | | | | |
| *Slc38a2* | 1.155 ± 0.033 | 1.414 ± 0.126 | 1.234 ± 0.111 | 1.361 ± 0.107 | P_Diet_ = 0.9026 |
|  |  |  |  |  | P_Sex_ = 0.0715 |
|  |  |  |  |  | P_Int_ = 0.5259 |
| *Slc2a1* | 1.618 ± 0.097 | 1.353 ± 0.068 | 1.442 ± 0.080 | 1.476 ± 0.121 | P_Diet_ = 0.7770 |
|  |  |  |  |  | P_Sex_ = 0.2216 |
|  |  |  |  |  | P_Int_ = 0.1168 |
| *Slc2a3* | 1.025 ± 0.064 | 1.015 ± 0.055 | 0.895 ± 0.081 | 1.124 ± 0.055 | P_Diet_ = 0.8671 |
|  |  |  |  |  | P_Sex_ = 0.0990 |
|  |  |  |  |  | P_Int_ = 0.0723 |
| *Fabp4* | 1.271 ± 0.101 | 1.514 ± 0.180 | 1.258 ± 0.077 | 1.348 ± 0.094 | P_Diet_ = 0.5587 |
|  |  |  |  |  | P_Sex_ = 0.2083 |
|  |  |  |  |  | P_Int_ = 0.05587 |
| **E18.5** | | | | | |
| *Slc38a2* | 0.725 ± 0.027^ab^ | 0.658 ± 0.051^a^ | 0.700 ± 0.067^ab^ | 0.860 ± 0.028^b^ | P_Diet_ = 0.0837 |
|  |  |  |  |  | P_Sex_ = 0.3475 |
|  |  |  |  |  | **P_Int_ = 0.0282** |
| *Slc2a1* | 0.762 ± 0.030 | 0.803 ± 0.065 | 0.738 ± 0.110 | 0.830 ± 0.033 | P_Diet_ = 0.9849 |
|  |  |  |  |  | P_Sex_ = 0.3222 |
|  |  |  |  |  | P_Int_ = 0.7030 |
| *Slc2a3* | 0.747 ± 0.026 | 1.060 ± 0.098 | 0.873 ± 0.109 | 0.904 ± 0.137 | P_Diet_ = 0.8685 |
|  |  |  |  |  | P_Sex_ = 0.0775 |
|  |  |  |  |  | P_Int_ = 0.1438 |
| *Fabp4* | 0.580 ± 0.066 | 0.727 ± 0.082 | 0.756 ± 0.050 | 0.848 ± 0.175 | P_Diet_ = 0.1413 |
|  |  |  |  |  | P_Sex_ = 0.2315 |
|  |  |  |  |  | P_Int_ = 0.7795 |

CON, placentae from litters generated by CON sires; PHF, placentae from litters generated by HF-fed sires; *Fabp4*, fatty acid binding protein 4; Int, interaction; *Slc2a1*, solute carrier family 2 member 1; *Slc2a3*, solute carrier family 2 member 3; *Slc38a2*, solute carrier family 38 member 2.

*Data are presented as mean ± SEM. Two-way ANOVA with main effects of sire diet and fetal sex as factors using Bonferroni’s post-hoc for multiple comparisons. Groups that do not share the same letter indicate significance *P* < 0.05.
