## Supplemental Table S8 for "Paternal obesity results in placental hypoxia and sex-specific impairments in placental vascularization and offspring metabolic function"

**Supplemental Table S8.** Maternal and fetal outcomes at embryonic day (E)14.5 and E18.5.*

| **E14.5** | | | | | |
| --- | --- | --- | --- | --- | --- |
| **Maternal/Litter Outcomes** | **Sire Group** | | | | **P-Value** |
|  | **CON**  (n = 33) | | **PHF**  (n = 24-25) | |  |
| Dam Gestational Mass Gained (g) | 5.0 ± 0.2 | | 4.9 ± 0.3 | | 0.9094 |
| Dam Fasting Blood Glucose (mmol/L) | 5.3 ± 0.2 | | 5.3 ± 0.2 | | 0.8384 |
| Litter Size (Fetuses/Litter) | 7.5 ± 0.2 | | 7.6 ± 0.3 | | 0.6739 |
| Number of Resorptions (Resorptions/Litter) | 1.5 ± 0.2 | | 0.9 ± 0.2 | | 0.1544 |
| Fetal Sex Ratio (Male:Female) | 1.3 ± 0.2 | | 1.1 ± 0.5 | | 0.3766 |
| **Fetal Outcome** | **Male**  (n = 10-28) | **Female**  (n = 11-29) | **Male**  (n = 10-23) | **Female**  (n = 11-22) | **P-Value** |
| Fetal Body Mass (g) | 0.247 ± 0.005^a^ | 0.227 ± 0.004^b^ | 0.237 ± 0.004^ab^ | 0.230 ± 0.005^ab^ | 0.0079 |
| Placental Mass (g) | 0.097 ± 0.002 | 0.092 ± 0.002 | 0.093 ± 0.002 | 0.118 ± 0.032 | 0.5923 |
| **E18.5** | | | | | |
| **Maternal/Litter Outcomes** | **Sire Group** | | | | **P-Value** |
|  | **CON**  (n = 24-30) | | **PHF**  (n = 18-24) | |  |
| Gestational Mass Gained (g) | 15.1 ± 0.4 | | 13.7 ± 0.6 | | 0.1481 |
| Dam Fasting Blood Glucose (mmol/L) | 4.9 ± 0.2 | | 4.8 ± 0.2 | | 0.5251 |
| Litter Size (Fetuses/Litter) | 8.0 ± 0.3 | | 7.0 ± 0.5 | | 0.3322 |
| Number of Resorptions (Resorptions/Litter) | 0.6 ± 0.2 | | 1.7 ± 0.5 | | 0.1122 |
| Fetal Sex Ratio (Male:Female) | 1.8 ± 0.3 | | 2.0 ± 0.4 | | 0.5615 |
| **Fetal Outcome** | **Male**  (n = 30) | **Female**  (n = 30) | **Male**  (n = 23) | **Female**  (n = 23) | **P-Value** |
| Fetal Body Mass (g) | 1.09 ± 0.02 | 1.05 ± 0.02 | 1.09 ± 0.02 | 1.08 ± 0.02 | 0.6227 |
| Placental Mass (g) | 0.127 ± 0.006 | 0.116 ± 0.006 | 0.124 ± 0.007 | 0.120 ± 0.007 | 0.6033 |

CON, dams mated with CON males or pregnancies generated by CON males; PHF, dams mated with high fat diet-fed males or pregnancies generated by high fat diet-fed males.

*Data are presented as mean ± SEM.

^#^Nested t-test or nested one way ANOVA using Bonferonni’s post-hoc for multiple comparison. Groups that do not share the same letter indicate significance *P* < 0.05.
